# INO80 regulates chromatin accessibility to facilitate suppression of sex-linked gene expression during mouse spermatogenesis

**DOI:** 10.1101/2023.01.04.522761

**Authors:** Prabuddha Chakraborty, Terry Magnuson

## Abstract

The INO80 protein is the main catalytic subunit of the INO80-chromatin remodeling complex, which is critical for DNA repair and transcription regulation in murine spermatocytes. In this study, we explored the role of INO80 in silencing genes on meiotic sex chromosomes in male mice. INO80 immunolocalization at the XY body in pachytene spermatocytes suggested a role for INO80 in the meiotic sex body. Subsequent deletion of *Ino80* resulted in high expression of sex-linked genes. Furthermore, the active form of RNA polymerase II at the sex chromosomes of *Ino80*-null pachytene spermatocytes indicates incomplete inactivation of sex-linked genes. A reduction in the recruitment of initiators of meiotic sex chromosome inhibition (MSCI) argues for INO80-facilitated recruitment of DNA repair factors required for silencing sex-linked genes. This role of INO80 is independent of a common INO80 target H2A.Z. Instead, in the absence of INO80, a reduction in chromatin accessibility at DNA repair sites occurs on the sex chromosomes. These data suggest a role for INO80 in DNA repair factor localization, thereby facilitating the silencing of sex-linked genes during the onset of pachynema.

**Summary Statement:** Chromatin accessibility and DNA repair factor localization at the sex chromosomes are facilitated by INO80, which regulates sex-linked gene silencing during meiotic progression in spermatocytes.

## Introduction

Mammalian gametogenesis involves the meiotic division of diploid germ cells that undergo homologous chromosome synapsis and recombination to generate haploid gametes. Extended prophase I during meiosis ensures the faithful execution of recombination to shuffle genetic material between homologous chromosomes by controlled introduction of DNA double-strand breaks (DSB) at the DSB hotspots by SPO11 followed by their repair (1). Synapsis between homologous autosomes occurs during the zygotene stage of meiotic prophase I. In contrast, non-homologous regions of the X and Y sex chromosomes in male germ cells do not synapse with each other (2).

During pachynema, DNA double-strand break repair (DSBR) factors no longer localize to autosomes, indicating the completion of DSBR. For the sex chromosomes, the DSBR factors sequester at unpaired regions, forming a unique chromatin domain called the sex body (3,4). The sex chromosomes in spermatocytes undergo modifications by several DNA damage repair (DDR) factors such as BRCA1, ATR, and its activator TOPBP1 at the unpaired (asynapsed) chromosome axes (5,6). The sex chromosomes also undergo chromatin remodeling to induce epigenetic silencing of sex-linked genes. This process is known as meiotic sex chromosome inactivation (MSCI) (7). Incomplete MSCI leads to germ cell death during pachynema, causing male infertility (7).

MSCI is initiated by DNA double-strand breaks (DSB) at the asynapsed axes and the incorporation of DSBR proteins at these DSB sites (8,9). ATR localizes to the DSBs along the chromosome axes (10). Serine-139 phosphorylation of H2A.X (γH2A.X) at the DSBs by ATR leads to the recruitment of another DSB factor, MDC1 (6,10–12). MDC1 localization amplifies γH2A.X in the protruding chromatin loops along the axes, establishing the characteristic sex-body and MSCI (6,13).

Chromatin remodelers regulate chromatin accessibility. They play roles in several cellular processes, including DNA repair and transcription regulation. Chromatin remodeling enzymes can hydrolyze ATP to change chromatin conformation by moving, evicting, or incorporating nucleosomes. They can also facilitate specific histone variant exchange to change local chromatin accessibility. There are four major families of chromatin remodeling complexes, all of which play critical functions in murine spermatogenesis (14–20). INO80 is the main ATPase subunit of INO80 complex, which has broad effects on several cellular processes. INO80 is important in a wide variety of DNA metabolic processes, including DNA replication, transcription, repair, and genome stability in several organisms (21). The exchange and turnover of histone variant H2A.Z is regulated by INO80 (22–25). INO80 facilitates the recruitment of RNA polymerase II (RNAPII) to the promoters of pluripotency network genes in ES cells (26,27). In yeast, INO80 is also implicated in the removal and degradation of ubiquitinated RNAPII from chromatin (28). Like other chromatin remodelers, INO80 is expressed in several mammalian tissues, including the testis (29), and was reported to facilitate development in a context-dependent manner (27,29–31). However, it is unclear whether chromatin remodeler INO80 plays any role in the meiotic silencing of the sex chromosomes.

Here, we explore the role of INO80 in silencing meiotic sex chromosomes. We show that INO80 interacts with and regulates sex-linked gene silencing in pachytene spermatocytes. Further, INO80 promotes the opening of the sex chromatin during the zygonema-to-pachynema transition at the DSB regions. INO80 also facilitates the recruitment of DNA repair factors at the DSB sites and interacts with DSBR factors to facilitate the silencing of sex chromosomes.

## Results

### INO80 localizes to the sex chromosomes in meiotic germ cells

Immunofluorescence localization of INO80 in pachytene spermatocytes revealed staining on the sex chromosomes (Fig. 1A, B). Chromatin immunoprecipitation followed by high throughput sequencing (ChIP-seq) for INO80 (GEO Dataset GSE179584) (32) also showed binding on the sex chromosomes (Fig. 1C). INO80 binding occurred at the zygotene and pachytene stages (GEO Dataset GSE190590) (33) on both autosomes and sex chromosomes. However, localization is more enriched on the sex chromosomes, especially during postnatal day 18 (P18) pachytene stage (Fig. 1C). Additionally, INO80 binding sites on sex chromosomes were present at the DSB sites identified by the meiotic DSB marker γH2A.X (GEO dataset GSE75221) (34) (Fig. S1A). These binding sites include promoters, intergenic regions, and gene bodies (Fig. S1A). Moreover, INO80 and γH2A.X binding in the sex chromosomes were correlated (Fig. S1B), where INO80 binding is enriched at the γH2A.X binding sites (Fig. S1C). These data suggested a possible role of INO80 in DSB repair and silencing the expression of genes on the sex chromosomes.

**Figure 1:**
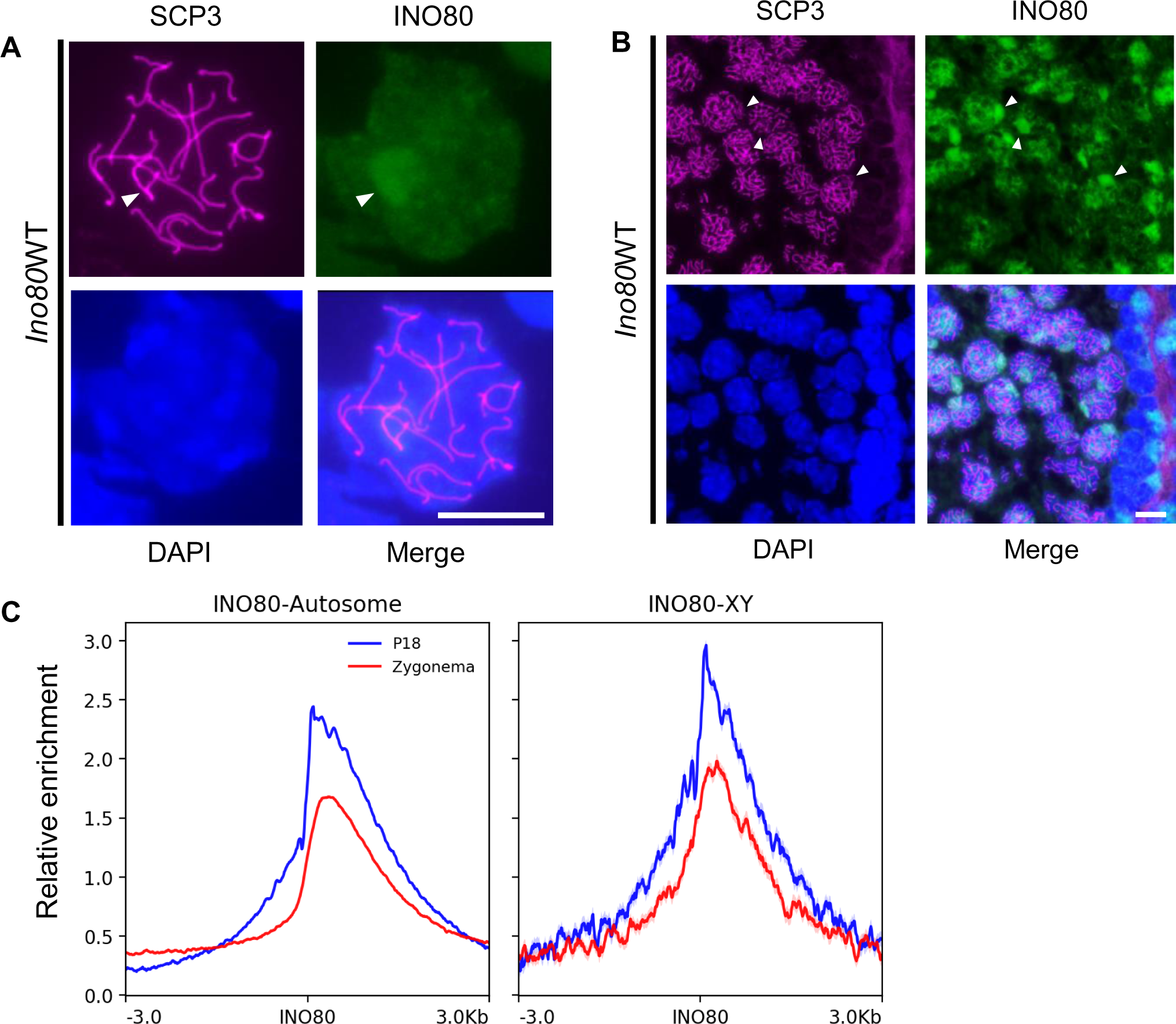
INO80 localization in the germ cells. (A,B) INO80 immunolocalization in the pachytene spermatocyte spreads (A) and the seminiferous tubule of the testis (B). SCP3 staining indicates the stage of the meiotic spermatocyte. Magenta: SCP3, Green: INO80, Blue: DAPI. Arrowhead: Sex chromosomes. Scale bar = 10µM. C; Enrichment of INO80 obtained by ChIP-seq on the autosomes and XY chromosomes during zygonema and at postnatal day 18 (P18), where most cells are in the pachytene stage. Blue: P18, Red: Zygonema. (Analyzed from GEO Dataset GSE179584) (32)

### INO80 regulates sex chromosome silencing in pachytene spermatocytes

RNAseq analysis of changes in P18 transcription from autosomes in wild type (*Ino80*WT) and germ cell-specific *Ino80*-null (*Ino80*cKO) spermatocytes (GEO Dataset GSE179584) (32) revealed up-and down-regulated genes (padj< 0.05). However, the mean changes in expression remained close to zero (Fig. 2A). In contrast, the differentially expressed genes (DEGs) (padj< 0.05) from the X- and Y-chromosomes remained upregulated in *Ino80*cKO cells (Fig. 2A). When plotted along the length of three representative autosomes and the sex chromosomes, like autosomal genes, DEGs from sex chromosomes occur along the length of the chromosomes (Fig. 2B). These data confirm that the lack of silencing of sex-linked genes is neither limited to a specific region such as the pseudo-autosomal region and not due to any positional effect. Quantitative RT-PCR analysis validated the upregulation of five representative sex-linked genes (*Ccnb3, Nxt2, Eda2r, Abcd1, Usp11*) in *Ino80*cKO spermatocytes, corroborating the RNAseq data. To investigate further the sex-linked gene expression in pachytene spermatocytes alone, we utilized meiotic germ cell synchronization to isolate homogeneous pachytene spermatocyte population (35) from wild-type and mutant testes and quantified sex-linked gene expression by qRT-PCR. We observed a similar pachytene spermatocyte population in both wild-type and mutant testes (Fig. S2A). Synchronized *Ino80*cKO pachytene spermatocytes also exhibited undetectable INO80 levels (Fig. S2B). An upregulation of all five genes (*Ccnb3, Nxt2, Eda2r, Abcd1, Usp11*) in the synchronized *Ino80*cKO pachytene spermatocytes confirms the lack of suppression of X-linked gene expression during pachynema upon *Ino80* deletion (Fig. S2C). Consistent with transcriptional silencing, immunofluorescence analysis of the active form of RNA polymerase II (pSer2) revealed its absence at the sex chromosomes in *Ino80*WT pachytene spermatocytes (Fig. 2D-E). These data correlate with normal transcriptional silencing. In contrast, continued RNA polymerase II (pSer2) immunofluorescence in *Ino80*cKO pachytene spermatocytes indicates incomplete silencing of the sex-linked genes (Fig. 2F-G). Overall, there was a significant increase in RNA polymerase II (pSer2) at the sex chromosomes in pachytene spermatocytes in the absence of INO80 (Fig. 2H).

**Figure 2:**
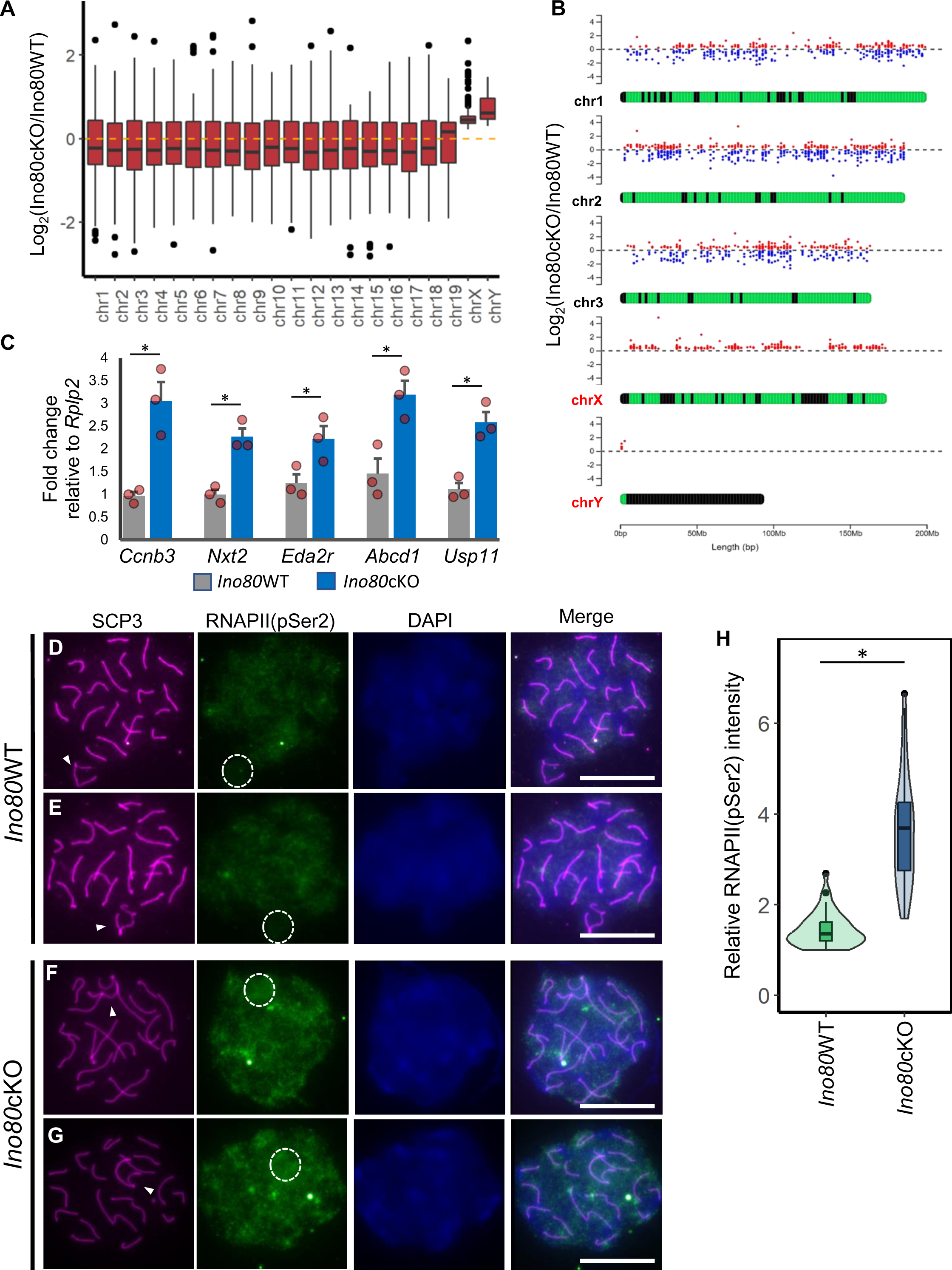
INO80 required for transcriptional silencing of sex chromosomes during meiotic prophase. (A) Boxplots showing mean differential gene expression from each chromosome in response to *Ino80* deletion in spermatocytes. The yellow dotted line indicates mean Log_2_FoldChange = 0. Chr: Chromosome. (n=5) (Analyzed from GEO Dataset GSE179584) (32) (B) Location of the differentially regulated genes along the length of three representative autosomes and the sex chromosomes. Dots above each chromosome indicate the fold change of individual genes. Upregulated and downregulated genes are indicated by red and blue, respectively. (Analyzed from GEO Dataset GSE179584) (32) (n=5). (C) Quantitative RT-PCR analysis of representative sex-linked gene expression levels normalized to *Rplp2* in either *Ino80*WT or *Ino80*cKO testes. Bars represent mean ± s.e.m. *; p<0.05, as calculated by unpaired t-test (n=3). (D-G) Immunolocalization of SCP3 (magenta) and RNA Polymerase II (pSer2) (green) in spermatocytes either from *Ino80*WT (D,E) or *Ino80*cKO (F,G) (Scale bar = 10µM) showing aberrant localization RNA Polymerase II (pSer2) (RNAPII) in *Ino80*cKO spermatocytes that are indicative of incomplete sex-chromosome silencing. White arrowhead; sex chromosome. The white circle denotes an approximate area of the sex chromosomes. (H) Relative quantification of the immunofluorescence signal intensity of RNAPII quantified from *Ino80*WT (n=50) and *Ino80*cKO (n=50) spermatocytes from three biological replicates. *; p<0.05, as calculated by Wilcoxon rank sum test.

### INO80 is necessary for the localization of DSBR factors to sex chromosomes

DSBR factors such as ATR, γH2A.X, and MDC1 are required to initiate meiotic sex chromosome inactivation in pachytene spermatocytes (6,10,36). We explored the localization of these DSBR proteins in P21 wild-type and mutant pachytene spermatocytes. Complete sequestration of ATR occurred on early *Ino80*WT pachytene sex chromosomes (Fig. 3A,C), where relatively consistent ATR binding occurs along the axis at the unsynapsed part of sex chromosomes (Fig. 3B,D). In contrast, ATR lacked uniform localization on the unsynapsed part of sex chromosomes of early to mid-pachytene *Ino80*cKO spermatocytes (Fig. 3E,G). Instead, upon quantitation, patchy ATR localization was observed along the axis (Fig. 3F, H). Further, the overall intensity of ATR at the sex chromosomes was moderately reduced (p<0.05) (Fig. 3J). Next, we tested the activity of ATR by immunostaining for the ATR substrate phospho-CHK1 (S345) (37). Although the localization of pCHK1 (S345) was limited to the sex chromosomes in *Ino80*WT pachytene spermatocytes, aberrant localization occurred in *Ino80*cKO spermatocytes (Fig. S3). These data indicate aberrant but active DNA binding and kinase activity of ATR in the *Ino80*cKO spermatocytes.

**Figure 3:**
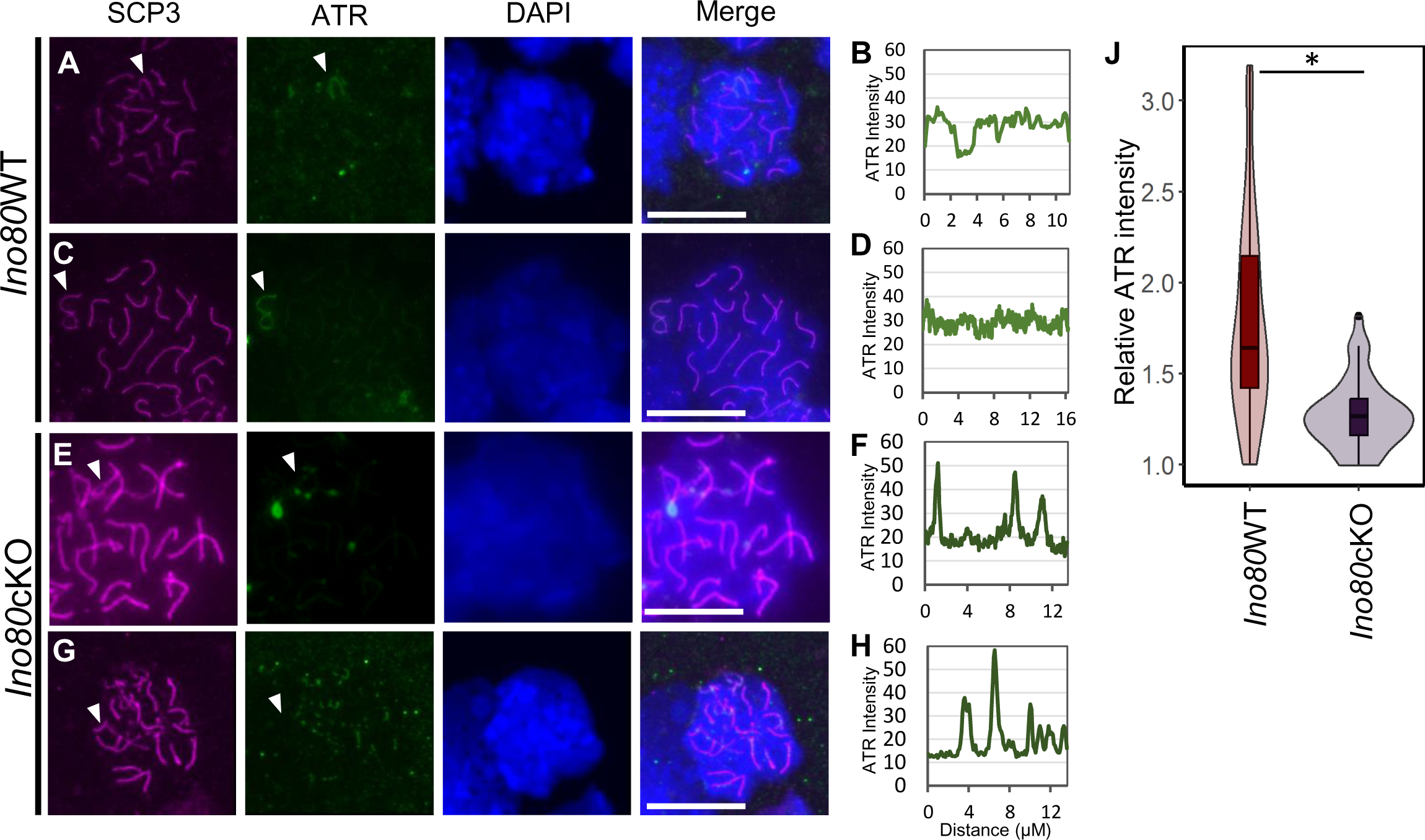
Aberrant ATR recruitment at sex chromosomes without INO80. (A-H) Immunolocalization of SCP3 (magenta) and ATR (green) in spermatocytes from *Ino80*WT (A,C) or *Ino80*cKO (E,G). DAPI is shown in blue. Scale bar = 10µM. White arrowhead; sex-chromosome. Line tracing for the quantification of ATR signal along the Y and X chromosome axes in the respective left panels from *Ino80*WT (B,D) and *Ino80*cKO (F,H) are displayed. (J) Relative fluorescent intensity measurement of ATR signal at the sex chromosomes from three biological replicates: *Ino80*WT (n=53) or *Ino80*cKO (n=42) pachytene spermatocytes. *; p<0.05, as calculated by Wilcoxon rank sum test.

During pachynema progression, the amplification of ATR-mediated phosphorylation of H2A.X depends on MDC1 recruitment (6). Next, we determined MDC1 localization in *Ino80*cKO pachytene spermatocytes. MDC1 immunofluorescence was visible on early-to late-pachynema *Ino80*WT sex chromosomes (Fig. 4A-C, G). Conversely, significantly reduced staining (p<0.05) occurred at the same stages in *Ino80*cKO spermatocytes (Fig. 4D-F, G). These results indicate that INO80 facilitates the localization of MDC1 at sex chromosome DSBs during pachynema. (23)

**Figure 4:**
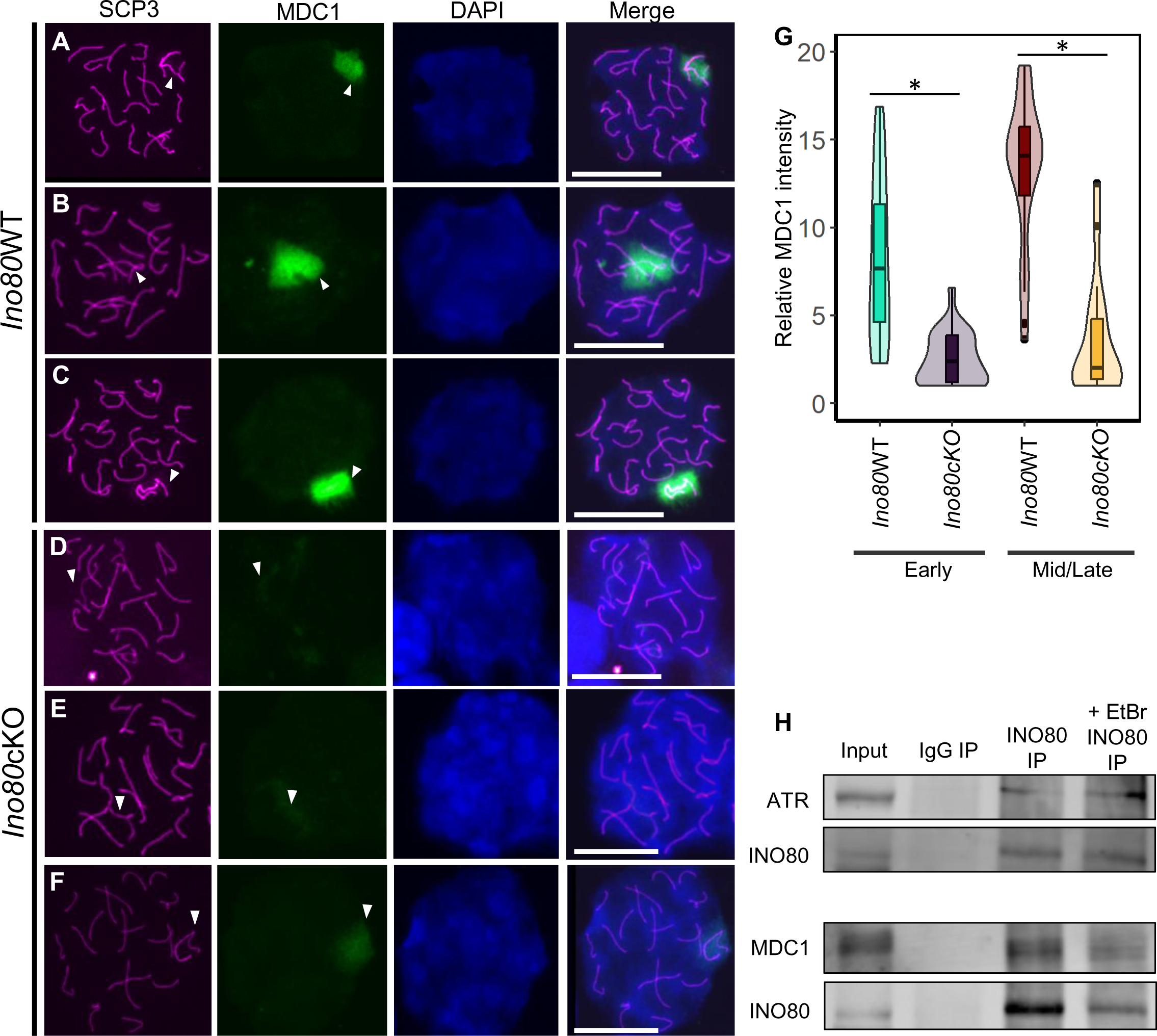
MDC1 recruitment at the sex chromosomes is facilitated by INO80. (A-F) Immunolocalization of SCP3 (magenta) and MDC1 (green) in spermatocytes from Ino80WT (A-C) or Ino80cKO (D-F). DAPI is shown in blue. Scale bar = 10µM. White arrowhead; sex-chromosome. (G) Relative fluorescent intensity measurement of MDC1 signal at the sex chromosomes from either *Ino80*WT (early; n=30, mid/late; n=39) or *Ino80*cKO (early; n=23, mid/late; n=17) pachytene spermatocytes from three biological replicates. *; p<0.05, as calculated by Wilcoxon rank sum test. (H) Immunoblot images demonstrating the interaction of INO80 with ATR and MDC1 by the presence of MDC1 and ATR in INO80 immunoprecipitated *Ino80*WT spermatocyte homogenates and INO80 when immunoprecipitated with MDC1 from *Ino80*WT spermatocyte homogenate.

To determine whether INO80 physically recruits DSB repair factors, we performed co-immunoprecipitation for INO80 from P21 spermatocytes. The presence of ATR and MDC1 with the INO80-immunoprecipitated samples (Fig. 4H) and the detection of INO80 in MDC1- and ATR-immunoprecipitated samples (Fig. S4A) supports the physical interaction between INO80 and DSBR factors ATR and MDC1. Adding either ethidium bromide or DNase to the spermatocyte lysate did not alter MDC1 or ATR detection in the immunoblot (Fig. 4H, S4B). This result indicates the interaction between MDC1 and INO80 is chromatin independent.

Further, to determine MDC1 recruitment, we performed <u>c</u>leavage <u>u</u>nder targets and release <u>u</u>sing <u>n</u>uclease (CUT&RUN) in synchronized *Ino80*WT and *Ino80*cKO pachytene spermatocytes. We observed robust MDC1 enrichment at INO80-binding sites only in the sex chromosomes in *Ino80*WT spermatocytes. In contrast, reduced MDC1 occupancy was present in Ino80cKO spermatocytes (Fig. 5A, S4C). Alternatively, in autosomes, MDC1 enrichment was absent (Fig. 5A). MDC1 occupancy was also substantially reduced at and around DSB sites marked by γH2A.X (Fig. 5B). Further, genomic annotation analysis of the MDC1 binding sites suggests majority of MDC1 occupancy was present in distal intergenic regions (63.95%). In contrast, reduced occupancy was distributed among promoters (9.19%), exons (6.08%), and introns (18.79%) (Fig. 5C). The reduction in MDC1 occupancy in *Ino80*cKO corroborates our previous observation by MDC1 immunofluorescence staining. Further, we also observed the downregulation of MDC1 mRNA and protein in the *Ino80*cKO spermatocytes (Fig. S4D,E). In addition, the enrichment of INO80 at the promoter of *Mdc1* suggests a requirement for INO80 regulation of MDC1 expression (Fig. S4F).

**Figure 5:**
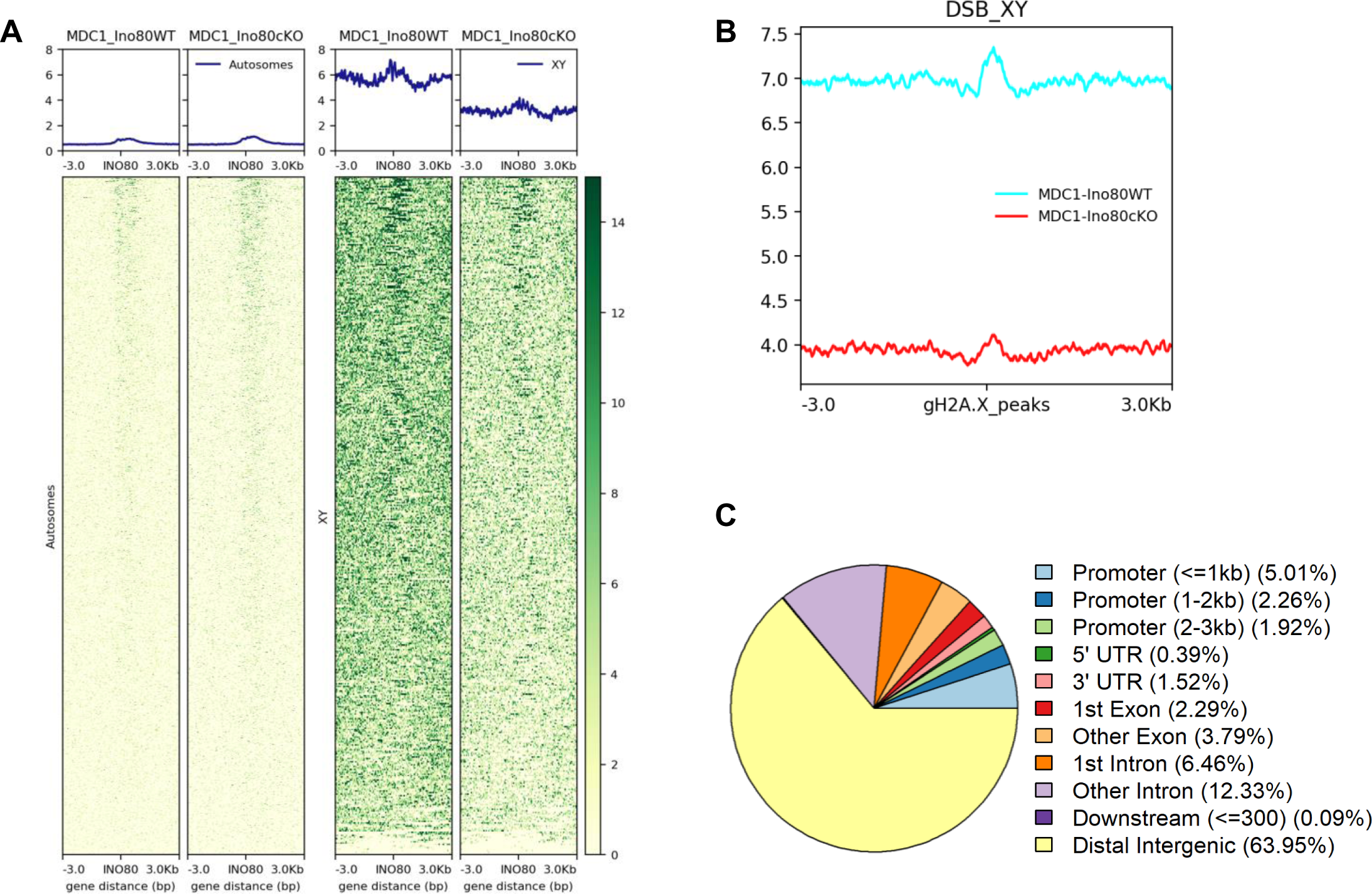
Genomic occupancy of MDC1. (A) Heatmap illustrating MDC1 occupancy at the INO80 binding sites in autosomes (left) and sex chromosomes (right). (B) Metaplot illustrating MDC1 occupancy at the DSB sites marked by γH2A.X binding. (C) Genomic annotation of MDC1 peaks in *Ino80*WT spermatocytes.

Because MDC1 is essential for amplifying γH2A.X, we determined whether perturbation of γH2A.X distribution occurred in the *Ino80*cKO spermatocytes. γH2A.X staining was observed in the *Ino80*WT pachytene spermatocytes (Fig. 6A,C), spanning the entire sex body (Fig. 6B,D). While γH2A.X staining was observed in the *Ino80*cKO spermatocytes (Fig. 6E,G), its expansion throughout the sex chromosomes seems perturbed and more concentrated near the axis (Fig. 6F,H). In addition, a moderate reduction in the overall γH2A.X signal at the sex chromosomes occurred in *Ino80*cKO spermatocytes, suggesting a perturbed amplification of γH2A.X (Fig. 6J).

**Figure 6:**
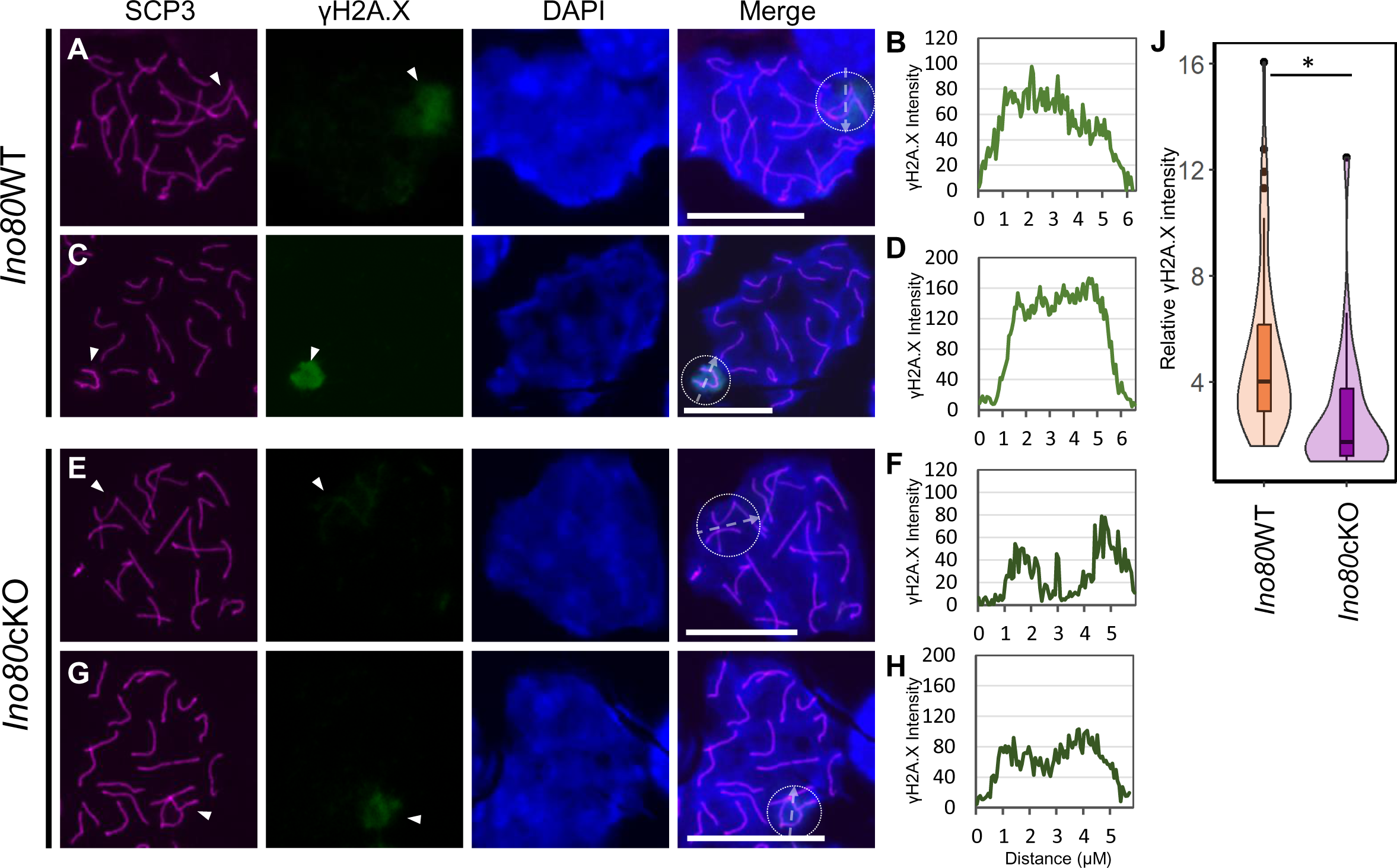
Aberrant. γ**H2A.X localization in *Ino80cKO* spermatocytes**. (A-H) Immunolocalization of SCP3 (magenta) and γH2A.X (green) in pachytene spermatocytes from *Ino80*WT (A,C) or *Ino80*cKO (E,G) (Scale bar = 10µM). DAPI is shown in blue. Scale bar = 10µM. White arrowhead; sex-chromosome. Line tracing for the quantification of γH2A.X signal along the dotted arrow in the dotted circle (marking the approximate sex body area) in the respective left panels from *Ino80*WT (B,D) and *Ino80*cKO (F,H) are displayed. (J) Relative fluorescent intensity measurement of γH2A.X signal at the sex chromosomes from either *Ino80*WT (n=58) or *Ino80*cKO (n=39) pachytene spermatocytes from three biological replicates. *; p<0.05, as calculated by Wilcoxon rank sum test.

### INO80 regulates chromatin accessibility at the sex chromosomes

INO80 is capable of histone exchange, most notably removing H2A.Z from chromatin (38,39). To determine how INO80 changes chromatin dynamics on the sex chromosomes and whether its regulation of sex-linked genes is H2A.Z-dependent, we compared H2A.Z occupancy (GEO Dataset GSE179584) (32) at sex-linked gene promoters in P18 *Ino80*cKO and *Ino80*WT spermatocytes. There was little change in H2A.Z levels at the promoter and transcriptional start sites (TSS) and INO80 binding sites on the sex chromosomes (Fig. S5A,B). These results suggest that H2A.Z does not mediate the sex-linked gene regulation by INO80. We also examined the occupancy of activating histone modification H3K4me3 and suppressive histone modification H3K27me3 at the sex-linked promoters. Neither showed any change (Fig. S5C,D) and did not mediate INO80-dependent gene regulation in the sex body.

Chromatin accessibility can be another regulator of DNA-interacting protein binding at the chromatin. A global change in chromatin accessibility occurs in developing spermatocytes during the mitosis-to-meiosis transition and subsequent progression through meiosis (40). We performed Assay for Transposase-Accessible Chromatin with high-throughput sequencing (ATAC-seq) to determine accessible chromatin distribution in the developing *Ino80*WT spermatocytes at P12 (enriched at zygonema) and compared it with ATAC-seq data from P18 *Ino80*WT spermatocytes (enriched at pachynema) (GEO Dataset GSE179584) (32). Chromatin accessibility at promoter/TSS regions in autosomes at both P12 and P18 was higher than that observed for the sex chromosomes (Fig. S6A). Further, an increase in chromatin accessibility occurred at these regions in both autosomes and sex chromosomes from P12 to P18 (Fig. S6A). This increase did not reverse in *Ino80*cKO spermatocytes at P18 (Fig. S6A). These data indicate a lack of requirement for INO80 in generating accessible chromatin at promoter/TSS during this transition. Next, we determined how the distribution of chromatin accessibility changed during the zygonema to pachynema transition. Comparison of sex chromosome-specific ATAC-peaks revealed a small portion of unique sites (1.34%) on P12 spermatocytes. Most P12 peaks are present at P18. The common peaks account for 20% of the P18 ATAC peaks from sex chromosomes, while 80% of ATAC peaks at P18 were *de novo* in nature (Fig. S6B). Comparison of ATAC-peaks on sex chromosomes from *Ino80*cKO and *Ino80*WT spermatocytes (GEO Dataset GSE179584) (32) showed only 17% of these *de novo* peaks remain in *Ino80*cKO, and 83% of de novo peaks are lost (Fig. S6B). Overall, the increase in chromatin accessibility across sex-chromosomes from P12 to P18 was reduced substantially both at all accessible sites (Fig. S6C) as well as INO80 binding sites (Fig. S6D). Further, genomic annotation analysis of these ATAC-peaks from sex chromosomes suggested a significant increase in intronic (9% to 19%) and distal intergenic (24% to 51%) accessible regions during the transition from P12 to P18 (Fig. S6E). This result is because 81% of the *de novo* gained peaks occurred in intronic and distal intergenic regions, and only 7% were at promoter/TSS sites (Fig. S6E). In *Ino80*cKO spermatocytes, we observed a loss of a similar distribution of peaks. Most of the binding occurred in intronic and intergenic regions, suggesting that INO80 regulates the generation of these peaks (Fig. S6E).

Next, we performed a chromosome-specific comparison of accessible regions between P12 and P18 spermatocytes from Ino80WT testes at each peak. These experiments revealed a significant increase in chromatin accessibility at various locations across autosomes and sex chromosomes (Fig. 7A). All autosomes showed an increase in chromatin accessibility at most of the differentially accessible (DA) regions (FDR <0.05) (Fig. 7A). However, sex chromosomes, especially the X-chromosome, exhibited a greater degree of increase at all the DA regions (FDR < 0.05) (Fig. 7A). The genomic locations of the DA sites mainly occur at the intronic and intergenic areas. A minor portion of them occur at promoter-proximal regions (Fig. 7B). The distribution of increased and decreased accessible areas (FDR < 0.05) was similar. In contrast, the number of regions with reduced accessibility was minimal. These data indicate that a genome-wide increase in chromatin accessibility occurs during the zygonema to pachynema transition. The increase is more prevalent on the sex chromosomes (Fig. 7A).

**Figure 7:**
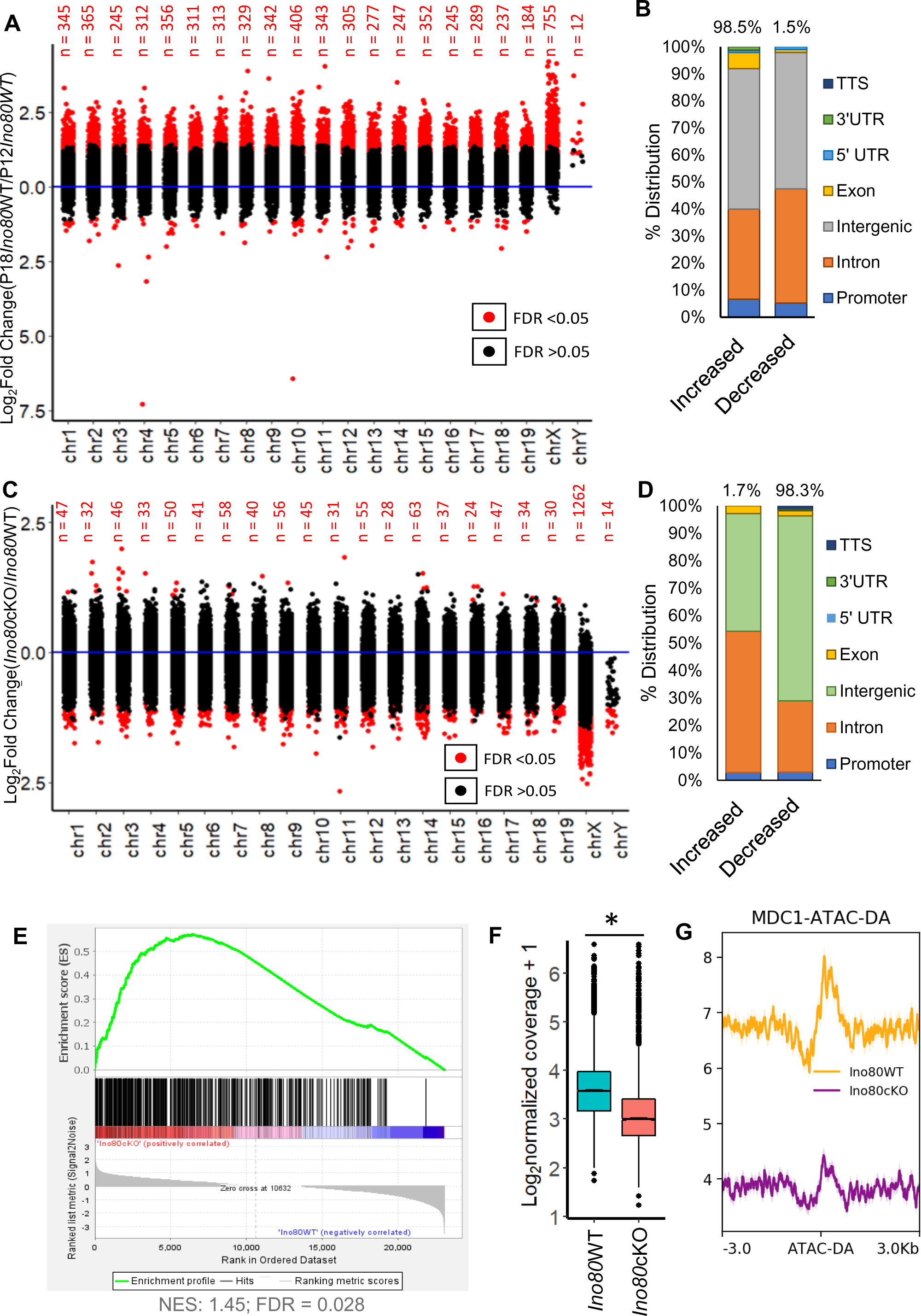
INO80 regulates chromatin accessibility in spermatocyte sex chromosomes. (A) Dot plot showing the relative changes in accessibility on each chromosome at P18 pachytene spermatocytes (GSE179584) (32) compared to P12 zygotene spermatocytes. Red dot: FDR < 0.05; Black dot: FDR > 0.05. FDR was derived by Benjamini-Hochberg method (n=3) (B) Genomic annotation of the differentially accessible regions from (FDR < 0.05) as shown in (A). (C) Dot plot showing relative changes in chromatin accessibility on each accessible site in each chromosome due to *Ino80* deletion at P18. Red dot: FDR < 0.05; Black dot: FDR > 0.05. FDR was derived by Benjamini-Hochberg method (n=3) (D) Genomic annotation of the differentially accessible regions from (FDR < 0.05) as shown in (C). (D); Genomic annotation of the differentially accessible regions due to *Ino80* deletion. (E) Gene set enrichment analysis with the nearest sex-linked genes of the differentially accessible regions at sex chromosomes in *Ino80*cKO spermatocytes, depicting enrichment of these genes among the upregulated genes in *Ino80*cKO, determined by RNAseq. NES; Normalized enrichment score. FDR; False discovery rate. (F) Boxplot showing the mean change in ATAC-signal (chromatin accessibility) at all the γH2A.X marked DSB sites (34) between *Ino80*WT and *Ino80*cKO in P18 spermatocytes. *; p<0.05, as calculated by Wilcoxon signed-rank test. (n=3). (G) Metaplot illustrating the reduction in the MDC1 occupancy at the differentially accessible regions in *Ino80*cKO sex chromosomes.

To determine the role of INO80 in regulating the transition in chromatin accessibility during meiotic progression, we compared the accessible sites in P18 spermatocytes from either *Ino80*WT or *Ino80*cKO spermatocytes (GEO Dataset GSE179584) (32). We observed a minimal change in accessible chromatin at the autosomes. In contrast, most of the DA regions were located at the sex chromosomes, demonstrating a much larger and significant change in accessibility (Fig. 7C). Genomic annotation analyses suggested that these DA chromatin regions mainly belong to intronic and intergenic regions (Fig. 7D). To determine how these DA sites were correlated to the change in transcription activity at the sex chromosomes, comparison of the nearest genes from the DA sites to the RNAseq data suggested significant enrichment of these sex-linked genes among the upregulated genes in *Ino80*cKO (FDR <0.05) (Fig.7E). These data indicate that INO80 plays a vital role in regulating the increased chromatin accessibility on the sex chromosome during meiotic progression. We next compared ATAC-signal from *Ino80*WT and *Ino80*cKO spermatocytes at the DSB regions marked by γH2A.X binding (obtained from publicly available dataset GSE75221) (34). A significant decrease in chromatin accessibility in *Ino80cKO* suggests a less permissive environment for DSBR factor recruitment at the DSB sites (Fig. 7F). We have also observed a decreased enrichment of MDC1 at the DA regions in *Ino80*cKO (Fig. 7G). These data indicate an essential role for INO80 in recruiting DNA damage repair factors to sex chromosome DSBs by regulating chromatin accessibility.

## Discussion

In this study, we demonstrated a unique role for INO80 in silencing sex-linked genes during pachynema in spermatocytes. INO80 regulates chromatin accessibility at DSB regions of the sex chromosomes. This regulation is independent of the histone variant H2A.Z, a major INO80 effector. We also demonstrated that INO80 interacts with DSBR factors ATR and MDC1, facilitating MDC1 recruitment at the sex chromosomes during pachynema.

An earlier report showed that male germ cell-specific deletion of *Ino80* results in a meiotic arrest phenotype in 8-week-old murine testes. A significant population of pachytene spermatocytes display defects in synapsis and DNA damage response, leading to the loss of spermatocytes in adult testes (18). However, during the first wave of spermatogenesis in juvenile mice, no significant cell death was observed in *Ino80*cKO testis up to P21. Moreover, unaltered gene expression signatures for zygonema and pachynema stages in *Ino80*cKO spermatocytes at P18, derived from RNAseq data comparing *Ino80*WT and *Ino80*cKo spermatocytes, suggest their relative proportions remain similar, except for a reduced transition from pachynema to diplonema in the absence of INO80 (32). Utilizing this small window of time, we observed a lack of downregulation of sex-linked differentially expressed genes in *Ino80*cKO spermatocytes, which was also replicated by comparing homogeneous pachynema populations upon synchronization of spermatogenesis. We also validated this overall aberrant sex-linked transcription program in individual *Ino80*cKO pachytene spermatocytes by immunolocalizing active RNA polymerase II at the sex chromosomes. This corroborates the observation of an incomplete MSCI in pachytene *Ino80*cKO spermatocytes. Several other studies also reported similar characteristics that described roles for MSCI-regulating factors (41–44).

Several studies have described a regulatory role for INO80 in DNA damage response in somatic cells and meiosis (18,45,46). The INO80 chromatin remodeling complex interacts with DNA damage factors such as γH2A.X and MEC1 (ATR) at DSB repair sites in yeast (47,48). We also found that INO80 interacts with DNA repair factors ATR and MDC1 in meiotic spermatocytes. γH2A.X recruitment at DSB sites during zygonema remained intact in *Ino80*cKO spermatocytes (18). We propose that the initial deposition of γH2A.X at the synapse axis of sex chromosomes initiates INO80-facilitated recruitment of MDC1. This recruitment amplifies γH2A.X in chromatin loops in an INO80-facilitated ATR-dependent fashion.

INO80 is known to regulate transcription in several cell types. It facilitates the recruitment of RNA polymerase II (RNAPII) and its cofactors to the promoters of pluripotency network genes. This recruitment regulates embryonic stem cell pluripotency and reprogramming (49,50). It can also regulate gene expression by facilitating histone modifications and the exchange of histone variants such as H2A.Z in different cell types (32,38,51,52). INO80 also regulates somatic gene silencing at the autosomes in spermatocytes by promoting H3K27me3 modification at promoters, while H3K4me3 remains unaffected (32). However, INO80-dependent silencing of sex-linked genes in meiotic spermatocytes is independent of INO80-mediated direct transcriptional regulation by enabling DNA binding factor recruitment at the promoter-proximal areas.

We showed that INO80 regulates sex chromosome accessibility during meiotic progression. Chromatin accessibility is a central regulator of DNA damage repair response. Less accessible DNA hinders successful and efficient DNA repair (53,54). Chromatin accessibility is also essential in transcription factor recruitment and efficient transcription regulation at the promoter-proximal areas (55). We observed an increase in overall chromatin accessibility at promoter regions of autosomes and sex chromosomes during meiotic progression.

In contrast, a similar level of accessibility remains at the promoter-proximal areas of sex chromosomes due to *Ino80 deletion*. Maezawa *et al.* previously reported that the accessibility at the promoter/TSS during the pachynema transition remains relatively unchanged. At the same time, gene expression changes occur (40). It is possible that chromatin accessibility is dynamic, and changes occur between stages during spermatogenesis. While the exact mechanism of gene silencing by DSB factors are not clear yet, the *de novo* generation of accessible regions at the non-promoter areas may provide the open chromatin structure necessary for DSB factors to bind and therefore explain resulting gene silencing by them during MSCI. Further, recent studies have reported a mechanistic view of the role of INO80 in sliding hexasomes and nucleosomes with different affinities to create accessible DNA (56,57). It is unclear whether hexasomes facilitate INO80-interaction at the meiotic sex chormosomes. However, it is logical to predict INO80 to be a central regulator of sex-linked gene silencing by regulating this *de novo* accessibility generation, as most of these peaks are lost in *Ino80*cKO spermatocytes. The loss of accessibility at the γH2A.X-binding DSB regions and reduced MDC1 occupancy at the DA sites in sex chromosomes due to *Ino80* deletion also indicates a possible INO80-dependent recruiting mechanism for DSBR factor MDC1.

Here we propose that INO80 mediates the *de novo* opening of chromatin around the DSB regions and facilitates recruitment of MDC1, whereby MDC1 initiates γH2A.X amplification by enabling ATR recruitment (Fig. 8).

**Figure 8:**
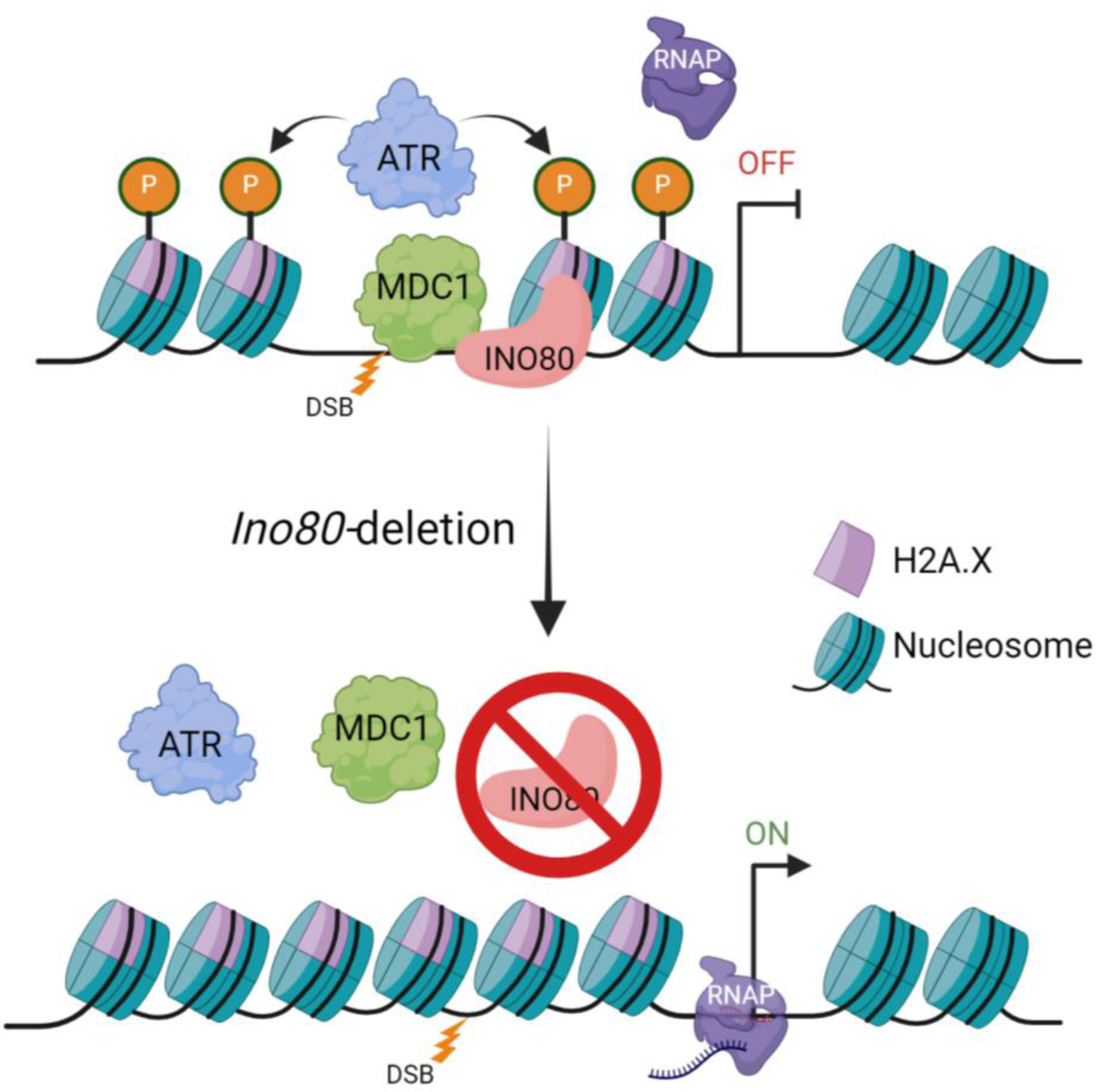
Schematic illustration of INO80-mediated regulation of meiotic sex-chromosome silencing. INO80 facilitates MDC1 recruitment at the DSB sites and regulates chromatin accessibility at these regions. MDC1 recruitment allows ATR-mediated amplification of γH2A.X at the chromatin loops. *Ino80*-deletion results in decreased chromatin accessibility and a lack of MDC1 recruitment, which fails to initiate ATR-mediated amplification of γH2A.X.

## Materials and methods

### Animals and genotyping

*Ino80* homozygous floxed (18,32) female mice (*Mus musculus*), maintained on an outbred CD1 background, were crossed with *Stra8-Cre^Tg/0^* males (58) to produce *Ino80^f/+^*; *Stra8Cre^Tg/0^* males. Male *Ino80^fl+^* (*Ino80*^WT^) and *Ino80*^Δ*/f*^*; Stra8Cre^Tg/0^* (*Ino80*c^KO^) littermates were obtained by crossing *Ino80^f/f^* females with *Ino80^f/+^; Stra8Cre^Tg/0^* males. Table S1 lists primers used for genotyping. Mice were maintained in an environment with controlled temperature and humidity with 12 h light and 12 h dark cycles and fed *ad libitum.* Spermatocyte synchronization was performed following a published protocol (35). In brief, newborn pups were treated with retinoic acid synthesis inhibitor WIN 18,446 (Cayman Chemical) (100µg/g), administered orally for 10 days (P1-P10). On P11, one bolus of retinoic acid (100ug per pup) was delivered subcutaneously to initiate spermatogonia differentiation, and pachytene spermatocytes were isolated on P24. All animal experiments were performed according to the protocol approved by the Institutional Animal Care and Use Committee at the University of North Carolina at Chapel Hill.

### RNA isolation and quantitative RT-PCR

Total RNA was isolated from Ino80WT and Ino80cKO spermatocytes using Trizol reagent (Invitrogen) followed by the Direct-zol RNA kit (Zymo). Reverse transcription was performed by the ProtoScript® II reverse transcriptase (NEB) using random primers. Real-time PCR was performed using Sso Fast EvaGreen supermix (Bio-Rad) on a thermocycler (Bio-Rad). Primers used in this study is listed in Table S2.

### Immunofluorescence staining

Spermatocyte nuclear spreads were prepared from freshly harvested testes following a published protocol (59) with modifications (60). Briefly, single-cell suspensions were prepared from seminiferous tubules. 3ml were added to three volumes of 0.25% NP40 (9ul) on a clean glass slide and incubated for 2 minutes at room temperature. Next, Fixative solution (36ul; 1% paraformaldehyde, 10 mM sodium borate buffer (pH 9.2) was added to the sample, and the slides were incubated in a moist chamber for 2 hours at room temperature. Lastly, slides were dried under a hood, washed with 0.5% Kodak Photo-Flo 200 three times, 1 minute each, and stored at -80°C.

Freshly harvested testes were embedded and frozen in the Optimum Cutting Temperature (OCT) embedding medium to make cryosections. Cryosections (7uM) on glass slides were fixed in freshly made 4% paraformaldehyde solution in PBS for 10 mins at 4°C. Following fixation, samples were washed in PBS 3 times, 5 minutes each, and permeabilized in PBST (PBS + 0.1% Triton-X 100) 3 times for 5 minutes each. Samples were incubated with blocking buffer (10% goat/donkey serum, 2% bovine serum albumin, 0.1% Triton-X 100 in PBS) for 1 hour at RT before incubation with primary antibody (listed in Table. S3) in blocking buffer overnight at 4°C. The following day, samples were washed 3 times, 5 minutes each with PBST, and incubated with Alexa Fluor-conjugated secondary antibody at room temperature for one hour. Samples were washed once with PBST and counterstained with DAPI, followed by three washes with PBST, 5 minutes each, and mounted in Prolong Gold anti-fade medium (P-36931; Life Technologies). Relative signal intensity was measured by NIH ImageJ software, either at an area marked by a region of interest with assigning the lowest intensity a value of 1 or along a single line transect as described previously (61). Meiotic substages were identified by either SCP1 and SCP3 immunostaining and/or the shape of the synapsed sex chromosomes (62). Plots were created in R using ggplot2 (63)

### Isolation of male germ cells

The spermatogenic cells were isolated following a modified version of a previously published protocol (64). Experiments were performed twice or more with cells isolated from separate mice. Briefly, freshly isolated testes were decapsulated, and seminiferous tubules were digested with collagenase (1mg/ml) in HBSS for 15 minutes at 32°C. Next, the tubules were precipitated by gravity for 5 mins at room temperature and separated, followed by a second digestion with collagenase (1 mg/ml) and trypsin (0.1%) in HBSS for 15 min at 32°C. Trypsin was inactivated by adding equal amounts of soybean trypsin inhibitor. The digested product was pipetted and filtered through 70 uM and then by 40uM cell strainers to get spermatocyte suspension. The spermatocytes were precipitated by centrifugation at 500x*g* for 10 mins, followed by two washes with HBSS. The cells were finally precipitated and either used in a following experiment or frozen at -80°C for later use.

### Nuclear lysate preparation

Nuclear lysate preparations occurred by isolating spermatogenic cells from P21 testes and incubating the cells in a hypotonic buffer (buffer A:10mM HEPES-KOH pH7.9, 1.5mM MgCl2, 10mM KCl, 0.1% NP-40, 5mM NaF, 1mM Na3VO4, 1mM PMSF, 1x Protease inhibitor cocktail) using 10-20 times the volume of the precipitated cell volume (PCV). After incubation on ice for 15 minutes, the cells were centrifuged at 1000x*g* for 10 minutes at 4°C. The cells were precipitated and resuspended in buffer A, using twice the PCV, and homogenized with a Dounce ‘B’ pestle 5 times on ice. Cells were precipitated by centrifugation at 1000 x *g* at 4°C. Precipitated nuclei were washed in buffer A and resuspended in equal volume lysis buffer (Buffer C) (20mM HEPES-KOH pH7.9, 1.5mM MgCl2, 420mM NaCl, 10mM KCl, 25 % glycerol, 0.2mM EDTA, 5mM NaF, 1mM Na3VO4, 1mM PMSF, 1x Protease inhibitor cocktail) for 30 minutes at 4°C on a nutator. The homogenate was cleared by centrifugation at 12000 x *g* for 10 mins at 4°C, and the supernatant saved in a separate tube. The extraction was performed again from the pellet, and the supernatant mixed with the previous one. The lysate was diluted with 2.8 volume of dilution buffer (Buffer D) (20mM HEPES-KOH pH7.9, 20 % glycerol, 0.2mM EDTA, 5mM NaF, 1mM Na3VO4, 1mM PMSF, 1x Protease inhibitor cocktail) to reduce the salt concentration and DTT added as necessary to a final concentration of 1mM. Some samples were treated with either ethidium bromide (50ug/ml) or DNase I (1µg/ml) to inhibit DNA-protein interaction (65) and followed by centrifugation at 12000 x *g* for 10 minutes at 4°C.

### Co-Immunoprecipitation

Co-immunoprecipitation was performed using 1-1.5 mg proteins from nuclear extracts from the spermatocytes. The lysate diluted with immunoprecipitation (IP) buffer (20mM HEPES-KOH pH7.9, 0.15mM KCl, 10% glycerol, 0.2mM EDTA, 0.5mM PMSF, 1x Protease inhibitor cocktail) to 1mg/ml concentration. For rabbit antibodies, Protein A conjugated Dynabeads (Invitrogen), and for mouse antibodies, Protein G conjugated Dynabeads (Invitrogen) were used (50ul per sample). Dynabeads were washed in PBS 3 times, 1 minute each, followed by incubation in PBS + 0.5% BSA for 10 minutes. Next, the beads were washed in PBS and IP buffer before use. Samples were precleared with dynabeads for 30 minutes at 4°C, followed by adding primary antibody (Table S3) and incubating at 4°C for 1 hour. Next, dynabeads were added to the samples and incubated overnight at 4°C. The next day, each sample was washed once in high salt wash buffer (20mM HEPES-KOH pH7.9, 300mM KCl, 10 % glycerol, 0.2mM EDTA, 0.1 % Tween-20, 1mM PMSF, Protease inhibitor cocktail), twice in IP wash buffer (20mM HEPES-KOH pH7.9, 150mM KCl, 10 % glycerol, 0.2mM EDTA, 0.1 % Tween-20, 1mM PMSF, Protease inhibitor cocktail), and once in final wash buffer (20mM HEPES-KOH pH7.9, 60mM KCl, 10 % glycerol, 1mM PMSF, 1x Protease inhibitor cocktail). Protein elution was performed using 1.3X Laemmli buffer and incubating at 65°C for 15 minutes, followed by magnetic removal of Dynabeads. The samples were finally heated at 95°C for 5 minutes to denature and stored at -20°C until used.

### Western Blotting

Proteins samples were subjected to polyacrylamide gel electrophoresis followed by overnight wet transfer to polyvinylidene difluoride (PVDF) membranes for fluorescence detection. Blots were blocked by Li-COR intercept blocking buffer followed by primary antibody (Table S3) incubation overnight in TBS with 0.1% Tween-20. Fluorescent Li-COR secondary antibodies were used for visualization, and blots were scanned using a Li-COR scanner. Uncropped blots are shown in the supplementary file 1.

### CUT&RUN

Cleavage under targets and release using nuclease (CUT&RUN) was performed using 250,000 spermatocytes per sample from Ino*80*WT and *Ino80*cKO testes following a previously published protocol (66). Briefly, a single cell suspension of spermatocytes was prepared and immediately washed three times, followed by attachment with concanavalin-A coated beads. These cells were permeabilized using digitonin and incubated overnight at 4°C with either IgG or antigen-specific primary antibody (Table S3). The next day, beads were washed twice, followed by protein-A/G-MNase binding in a calcium-free environment. Following two more washes to remove unbound Protein-A/G-MNase, calcium was introduced to start chromatin digestion for 30 mins at 0°C. Digested chromatin fragments were released for 30 mins at 37°C. The released chromatin was purified using DNA purification columns (Zymo ChIP DNA Clean & Concentrator). The elute was quantitated, followed by library preparation and high throughput sequencing by Novaseq X Plus.

### ATAC-seq

ATAC-seq was performed following the omni-ATAC method described previously (67), using *Ino80*WT spermatocytes from P12 testes. 50,000 cells from each sample were washed, and nuclei were isolated. A transposition reaction was performed with these nuclei at 37°C for 30 minutes in a thermomixer at 1000 r.p.m. followed by a clean-up step using Zymo DNA Clean and Concentrator-5 columns. Libraries were amplified, size selected using 0.5X and 1.8X Kapa pure beads to generate a size range of ∼150bp to ∼2kb and quantified using NEBNext® kit for Illumina. Libraries were pooled and sequenced on a Novaseq 6000 platform generating 50bp paired-end reads.

### Data analysis

ChIP-sequencing data and CUT&RUN data were trimmed as necessary using trimmomatic (68). The reads were aligned to mouse reference genome mm10 using Bowtie2 (69) with sensitive settings. Samtools (70) was used to de-duplicate and merge the alignments. Deeptools (71) was used to make depth-normalized coverage tracks and metaplots after removing the mm10 blacklisted regions. Correlation analysis between ChIP-seq datasets were also performed using deeptools multiBamSummary tool using mapping quality >30. MACS2 (72) was used to call peaks using the options - extsize set to 147 and -nomodel.

RNAseq reads were aligned by Tophat2 (73) to mm10, and read counts were obtained by HTseqCount (74). DESeq2 (75) was used with recommended settings for differential expression analysis. Plots were prepared by ggplot2 (63) and chromoMap (76). ATAC-seq reads were processed by nf-core/atacseq (ver 1.1.0) pipeline (77). Briefly, reads were trimmed by Trim Galore (https://www.bioinformatics.babraham.ac.uk/projects/trim_galore/) and aligned to mouse reference genome mm10 using BWA. Picard (https://broadinstitute.github.io/picard/) was used to mark the duplicates, and normalized coverage tracks scaled to 1 million mapped reads were prepared by BEDTools (https://bedtools.readthedocs.io/en/latest/index.html). Differential accessibility analysis was performed by CSAW (78) using region-based binned read count followed by TMM normalization using genome-wide background estimation. Genomic annotation of peaks was performed by Homer (79). Plots were created in R using ggplot2 (63).

### Statistical analysis

Comparison of signals from ATAC-seq, immunofluorescence and qPCR experiments were tested by Wilcoxon signed rank test for paired observations or Wilcoxon rank sum test or unpaired t-test for unpaired observations. All the tests performed were two tailed.

## Acknowledgment

We thank Magnuson lab members for their input and comments.

## Author contribution

P.C. and T.M. conceived the study; P.C. carried out experiments, and data were analyzed and interpreted by P.C. and T.M. The manuscript was written by P.C. and edited by T.M.

## Data availability

The sequencing data from RNA-seq, ChIP-seq, CUT&RUN, and ATAC-seq experiments are available in the GEO database (Accession numbers GSE179584 and GSE221682).

---

**Figure S1:** INO80 binding at the DSB sites. (A) Genomic tracks illustrating INO80 binding at the DSB sites marked by γH2A.X (34) in P18 (32) and zygotene (33) spermatocytes. Each separate genomic location is denoted by alternating background coloring. IG-Up; Intergenic-Upstream. (B) Correlation analysis of INO80 and γH2A.X binding at the X and Y chromosomes. The numbers in the box represent Pearson’s correlation coefficient calculated from high confidence reads (mapping quality >30) mapped to either chromosome X or Y. (C) Metaplot showing INO80 occupancy in *Ino80*WT spermatocytes at the sex chromosome DSB sites marked by γH2A.X.

**Figure S2:** Synchronized pachytene spermatocytes exhibit a similar lack of sex-linked gene expression. (A) Comparison of spermatocyte population in synchronized P24 *Ino80*WT and *Ino80*cKO testes. (B) Immunoblot for INO80 and alpha-actin in synchronized P24 *Ino80*WT and *Ino80*cKO testes. (C) Quantitative RT-PCR analysis of representative sex-linked gene expression levels normalized to *Rplp2* in synchronized P24 *Ino80*WT and *Ino80*cKO testes. Bars represent mean ± s.e.m. *; p<0.05, as calculated by unpaired t-test (n=3)

**Figure S3:** ATR substrate CHK1 phosphorylation remains intact without INO80. Immunolocalization of SCP3 (magenta) and pCHK1(S345) (green) in spermatocytes from Ino80WT or Ino80cKO. DAPI is shown in blue. Scale bar = 10µM. White arrowhead; sex-chromosome.

**Figure S4:** INO80 facilitates MDC1 recruitment and expression. (A) Immunoblot images demonstrate the interaction between INO80 with ATR and MDC1 by the presence of INO80 in both ATR and MDC1 immunoprecipitated samples. The top panel shows two brightness and contrast levels from the same blot to visualize MDC1 in input and immunoprecipitation samples. Spliced sections in the bottom panel are part of the same blot. (B) Immunoblot images demonstrate the presence of ATR and MDC1 in INO80-immunoprecipitated sample in the presence of DNase I. (C) Genomic tracks depicting the normalized enrichment of MDC1 at the representative sex-linked DEGs in *Ino80*WT and *Ino80*cKO pachytene spermatocytes. (D) Boxplot showing the normalized count of *Mdc1* transcripts from *Ino80*WT and *Ino80*cKO spermatocytes. (n=5) (Analyzed from GEO Dataset GSE179584) (32) (E) Immunoblot showing MDC1 (top) and α-Actin (bottom) expression from *Ino80*WT and *Ino80*cKO spermatocytes on P18. (F) Genomic tracks illustrated the enrichment of INO80, H3K27me3, and H3K4me3 at the *Mdc1* promoter-proximal area. (Analyzed from GEO Dataset GSE179584) (32)

**Figure S5:** Histone modifications at the sex chromosomes. Metaplot illustrating changes in H2A.Z (A-B), H3K4me3 (C), and H3K27me3 (D) occupancy at either INO80 binding sites (A) or promoter/TSS regions (B-D) in sex chromosomes (Analyzed from GEO Dataset GSE179584) (32).

**Figure S6:** Chromatin accessibility during meiotic progression. (A) Metaplot illustrating chromatin accessibility at the promoter/TSS regions of autosomes and sex chromosomes in either P12 *Ino80*WT or P18 *Ino80*WT and *Ino80*cKO spermatocytes. (B) Comparison of ATAC peaks at sex chromosomes during the transition from P12 spermatocyte to P18 spermatocyte and in P18 *Ino80*WT vs. *Ino80*cKO spermatocytes. (C-D) Metaplot illustrating chromatin accessibility in P12 *Ino80*WT and P18 *Ino80*WT and *Ino80*cKO spermatocytes at all the ATAC peaks (C) and at the INO80 peaks (D). (E) Genomic annotation of ATAC-peaks in P12 and P18 spermatocytes, as well as the *de novo* peaks generated during pachynema transition during P18 and lost due to *Ino80* deletion. P18 ATAC-seq data was analyzed from GEO Dataset GSE179584 (32).

## References

1. Handel MA, Schimenti JC. Genetics of mammalian meiosis: Regulation, dynamics and impact on fertility. Nat Rev Genet. 2010 Jan 29;11(2):124–36.

2. Burgoyne PS, Baker TG. Perinatal oocyte loss in XO mice and its implications for the aetiology of gonadal dysgenesis in XO women. J Reprod Fertil. 1985 Nov;75(2):633–45.

3. Solari AJ. The morphology and ultrastructure of the sex vesicle in the mouse. Exp Cell Res. 1964;36(1):160–8.

4. Yan W, McCarrey JR. Sex chromosome inactivation in the male. Vol. 4, Epigenetics. Taylor and Francis Inc.; 2009. p. 452–6.

5. Turner JMA, Aprelikova O, Xu X, Wang R, Kim S, Chandramouli GVR, et al. BRCA1, histone H2AX phosphorylation, and male meiotic sex chromosome inactivation. Current Biology. 2004 Dec 14;14(23):2135–42.

6. Ichijima Y, Ichijima M, Lou Z, Nussenzweig A, Daniel Camerini-Otero R, Chen J, et al. MDC1 directs chromosome-wide silencing of the sex chromosomes in male germ cells. Genes Dev. 2011 May 1;25(9):959–71.

7. Turner JMA. Meiotic Silencing in Mammals. Annu Rev Genet. 2015;49:395–412.

8. Carofiglio F, Inagaki A, de Vries S, Wassenaar E, Schoenmakers S, Vermeulen C, et al. SPO11-Independent DNA Repair Foci and Their Role in Meiotic Silencing. PLoS Genet. 2013 Jun;9(6).

9. Schoenmakers S, Wassenaar E, van Cappellen WA, Derijck AA, de Boer P, Laven JSE, et al. Increased frequency of asynapsis and associated meiotic silencing of heterologous chromatin in the presence of irradiation-induced extra DNA double strand breaks. Dev Biol. 2008 May 1;317(1):270–81.

10. Royo H, Prosser H, Ruzankina Y, Mahadevaiah SK, Cloutier JM, Baumann M, et al. ATR acts stage specifically to regulate multiple aspects of mammalian meiotic silencing. Genes Dev. 2013 Jul 1;27(13):1484–94.

11. Stewart GS, Wang B, Bigneli CR, Taylor AMR, Elledge SJ. MDC1 is a mediator of the mammalian DNA damage checkpoint. Nature. 2003 Feb 27;421(6926):961–6.

12. Stucki M, Clapperton JA, Mohammad D, Yaffe MB, Smerdon SJ, Jackson SP. MDC1 directly binds phosphorylated histone H2AX to regulate cellular responses to DNA double-strand breaks. Cell. 2005;123(7):1213–26.

13. Ichijima Y, Sin HS, Namekawa SH. Sex chromosome inactivation in germ cells: Emerging roles of DNA damage response pathways. Vol. 69, Cellular and Molecular Life Sciences. Springer; 2012. p. 2559–72.

14. Dowdle JA, Mehta M, Kass EM, Vuong BQ, Inagaki A, Egli D, et al. Mouse BAZ1A (ACF1) Is Dispensable for Double-Strand Break Repair but Is Essential for Averting Improper Gene Expression during Spermatogenesis. Schimenti JC, editor. PLoS Genet. 2013 Nov 7;9(11):e1003945.

15. Imai Y, Biot M, Clément JAJ, Teragaki M, Urbach S, Robert T, et al. Prdm9 activity depends on hells and promotes local 5-hydroxymethylcytosine enrichment. Elife. 2020;9:e57117.

16. Kim Y, Fedoriw AM, Magnuson T. An essential role for a mammalian SWI/SNF chromatin-remodeling complex during male meiosis. Development. 2012;139(6):1133–40.

17. Li W, Wu J, Kim SY, Zhao M, Hearn SA, Zhang MQ, et al. Chd5 orchestrates chromatin remodelling during sperm development. Nat Commun. 2014 May 13;5:3812.

18. Serber DW, Runge JS, Menon DU, Magnuson T. The mouse INO80 chromatin-remodeling complex is an essential meiotic factor for spermatogenesis. Biol Reprod. 2016;94(1):1–9.

19. Spruce C, Dlamini S, Ananda G, Bronkema N, Tian H, Paigen K, et al. HELLS and PRDM9 form a pioneer complex to open chromatin at meiotic recombination hot spots. Genes Dev. 2020;34(5):398–412.

20. Wang J, Gu H, Lin H, Chi T. Essential roles of the chromatin remodeling factor BRG1 in spermatogenesis in mice. Biol Reprod. 2012 Jun 22;86(6):186.

21. Conaway RC, Conaway JW. The INO80 chromatin remodeling complex in transcription, replication and repair. Trends Biochem Sci. 2009;34(2):71–7.

22. Alatwi HE, Downs JA. Removal of H2A.Z by INO 80 promotes homologous recombination. EMBO Rep. 2015;16(8):986–94.

23. Brahma S, Udugama MI, Kim J, Hada A, Bhardwaj SK, Hailu SG, et al. INO80 exchanges H2A.Z for H2A by translocating on DNA proximal to histone dimers. Nat Commun. 2017;8(15616):15616.

24. Papamichos-Chronakis M, Watanabe S, Rando OJ, Peterson CL. Global regulation of H2A.Z localization by the INO80 chromatin-remodeling enzyme is essential for genome integrity. Cell. 2011 Jan 21;144(2):200–13.

25. Ranjan A, Mizuguchi G, Fitzgerald PC, Wei D, Wang F, Huang Y, et al. Nucleosome-free region dominates histone acetylation in targeting SWR1 to promoters for H2A.Z replacement. Cell. 2013;154(6):1232–45.

26. Wang L, Du Y, Ward JM, Shimbo T, Lackford B, Zheng X, et al. INO80 facilitates pluripotency gene activation in embryonic stem cell self-renewal, reprogramming, and blastocyst development. Cell Stem Cell. 2014 May 1;14(5):575–91.

27. Zhou B, Wang L, Zhang S, Bennett BD, He F, Zhang Y, et al. INO80 governs superenhancer-mediated oncogenic transcription and tumor growth in melanoma. Genes Dev. 2016 Jun 15;30(12):1440–53.

28. Lafon A, Taranum S, Pietrocola F, Dingli F, Loew D, Brahma S, et al. INO80 Chromatin Remodeler Facilitates Release of RNA Polymerase II from Chromatin for Ubiquitin-Mediated Proteasomal Degradation. Mol Cell. 2015 Dec 3;60(5):784–96.

29. Serber DW, Runge JS, Menon DU, Magnuson T. The mouse INO80 chromatin-remodeling complex is an essential meiotic factor for spermatogenesis. Biol Reprod. 2016;94(1):1–9.

30. Ren Z, Zhao W, Li D, Yu P, Mao L, Zhao Q, et al. INO80-Dependent Remodeling of Transcriptional Regulatory Network Underlies the Progression of Heart Failure. Circulation. 2024 Apr 2;149(14):1121–38.

31. Gourisankar S, Krokhotin A, Wenderski W, Crabtree GR. Context-specific functions of chromatin remodellers in development and disease. Vol. 25, Nature Reviews Genetics. Nature Research; 2024. p. 340–61.

32. Chakraborty P, Magnuson T. INO80 requires a polycomb subunit to regulate the establishment of poised chromatin in murine spermatocytes. Development (Cambridge). 2022 Jan 1;149(1).

33. Carbajal A, Gryniuk I, Castro RO de, Pezza RJ. Efficient enrichment of synchronized mouse spermatocytes suitable for genome-wide analysis. bioRxiv. 2022 Jan 12;2022.01.11.475957.

34. Fayer S, Yu Q, Kim J, Moussette S, Camerini-Otero RD, Naumova AK. Robertsonian translocations modify genomic distribution of γH2AFX and H3.3 in mouse germ cells. Mammalian Genome. 2016;27(5–6):225–36.

35. Kirsanov O, Johnson T, Malachowski T, Niedenberger BA, Gilbert EA, Bhowmick D, et al. Modeling mammalian spermatogonial differentiation and meiotic initiation in vitro. Development (Cambridge). 2022 Nov 1;149(22).

36. Fernandez-Capetillo O, Mahadevaiah SK, Celeste A, Romanienko PJ, Camerini-Otero RD, Bonner WM, et al. H2AX is required for chromatin remodeling and inactivation of sex chromosomes in male mouse meiosis. Dev Cell. 2003 Apr 1;4(4):497–508.

37. Fedoriw AM, Menon D, Kim Y, Mu W, Magnuson T. Key mediators of somatic ATR signaling localize to unpaired chromosomes in spermatocytes. Development (Cambridge). 2015;142(17):2972–80.

38. Brahma S, Udugama MI, Kim J, Hada A, Bhardwaj SK, Hailu SG, et al. INO80 exchanges H2A.Z for H2A by translocating on DNA proximal to histone dimers. Nat Commun. 2017;8(15616):15616.

39. Papamichos-Chronakis M, Watanabe S, Rando OJ, Peterson CL. Global regulation of H2A.Z localization by the INO80 chromatin-remodeling enzyme is essential for genome integrity. Cell. 2011 Jan 21;144(2):200–13.

40. Maezawa S, Yukawa M, Alavattam KG, Barski A, Namekawa SH. Dynamic reorganization of open chromatin underlies diverse transcriptomes during spermatogenesis. Nucleic Acids Res. 2018;46(2):593–608.

41. Hirota T, Blakeley P, Sangrithi MN, Mahadevaiah SK, Encheva V, Snijders AP, et al. SETDB1 Links the Meiotic DNA Damage Response to Sex Chromosome Silencing in Mice. Dev Cell. 2018;47(5):645–659.e6.

42. Yeo AJ, Becherel OJ, Luff JE, Graham ME, Richard D, Lavin MF. Senataxin controls meiotic silencing through ATR activation and chromatin remodeling. Cell Discov. 2015 Sep 29;1:15025.

43. Becherel OJ, Yeo AJ, Stellati A, Heng EYH, Luff J, Suraweera AM, et al. Senataxin Plays an Essential Role with DNA Damage Response Proteins in Meiotic Recombination and Gene Silencing. McKinnon PJ, editor. PLoS Genet. 2013 Apr 11;9(4):e1003435.

44. Ichijima Y, Ichijima M, Lou Z, Nussenzweig A, Daniel Camerini-Otero R, Chen J, et al. MDC1 directs chromosome-wide silencing of the sex chromosomes in male germ cells. Genes Dev. 2011 May 1;25(9):959–71.

45. Morrison AJ. Genome maintenance functions of the INO80 chromatin remodeller. Philosophical Transactions of the Royal Society B: Biological Sciences. 2017;372(1731):20160289.

46. Poli J, Gasser SM, Papamichos-Chronakis M. The INO80 remodeller in transcription, replication and repair. Philosophical Transactions of the Royal Society B: Biological Sciences. 2017;372(1731):20160290.

47. Poli J, Gerhold CB, Tosi A, Hustedt N, Seeber A, Sack R, et al. Mec1, INO80, and the PAF1 complex cooperate to limit transcription replication conflicts through RNAPII removal during replication stress. Genes Dev. 2016;30(3):337–54.

48. Morrison AJ, Highland J, Krogan NJ, Arbel-Eden A, Greenblatt JF, Haber JE, et al. INO80 and γ-H2AX Interaction Links ATP-Dependent Chromatin Remodeling to DNA Damage Repair. Cell. 2004 Dec 17;119(6):767–75.

49. Wang L, Du Y, Ward JM, Shimbo T, Lackford B, Zheng X, et al. INO80 facilitates pluripotency gene activation in embryonic stem cell self-renewal, reprogramming, and blastocyst development. Cell Stem Cell. 2014 May 1;14(5):575–91.

50. Zhou B, Wang L, Zhang S, Bennett BD, He F, Zhang Y, et al. INO80 governs superenhancer-mediated oncogenic transcription and tumor growth in melanoma. Genes Dev. 2016 Jun 15;30(12):1440–53.

51. Yu H, Wang J, Lackford B, Bennett B, Li J liang, Hu G. INO80 promotes H2A.Z occupancy to regulate cell fate transition in pluripotent stem cells. Nucleic Acids Res. 2021;49(12):6739–55.

52. Alatwi HE, Downs JA. Removal of H2A.Z by INO 80 promotes homologous recombination. EMBO Rep. 2015 Aug 1;16(8):986–94.

53. Luijsterburg MS, Van Attikum H. Chromatin and the DNA damage response: The cancer connection. Vol. 5, Molecular Oncology. John Wiley & Sons, Ltd; 2011. p. 349–67.

54. Price BD, D’Andrea AD, Ahel D, Horejsí Z, Wiechens N, Polo SE, et al. Chromatin remodeling at DNA double-strand breaks. Cell. 2013 Mar 14;152(6):1344–54.

55. Allis CD, Jenuwein T. The molecular hallmarks of epigenetic control. Nature Reviews Genetics 2016 17:8. 2016 Jun 27;17(8):487–500.

56. Zhang M, Jungblut A, Kunert F, Hauptmann L, Hoffmann T, Kolesnikova O, et al. Hexasome-INO80 complex reveals structural basis of noncanonical nucleosome remodeling. Science (1979). 2023 Jul 21;381(6655):313–9.

57. Wu H, Muñoz EN, Hsieh LJ, Chio US, Gourdet MA, Narlikar GJ, et al. Reorientation of INO80 on hexasomes reveals basis for mechanistic versatility. Science (1979). 2023 Jul 21;381(6655):319–24.

58. Sadate-Ngatchou PI, Payne CJ, Dearth AT, Braun RE. Cre recombinase activity specific to postnatal, premeiotic male germ cells in transgenic mice. Genesis. 2008;46(12):738–42.

59. Wojtasz L, Daniel K, Roig I, Bolcun-Filas E, Xu H, Boonsanay V, et al. Mouse HORMAD1 and HORMAD2, two conserved meiotic chromosomal proteins, are depleted from synapsed chromosome axes with the help of TRIP13 AAA-ATPase. Lichten M, editor. PLoS Genet. 2009 Oct 23;5(10):e1000702.

60. Biswas U, Stevense M, Jessberger R. SMC1α Substitutes for Many Meiotic Functions of SMC1β but Cannot Protect Telomeres from Damage. Current Biology. 2018 Jan 22;28(2):249–261.e4.

61. Sin HS, Barski A, Zhang F, Kartashov A V., Nussenzweig A, Chen J, et al. RNF8 regulates active epigenetic modifications and escape gene activation from inactive sex chromosomes in post-meiotic spermatids. Genes Dev. 2012;26(24):2737–48.

62. Goetz P, Chandley AC, Speed RM. Morphological and temporal sequence of meiotic prophase development at puberty in the male mouse. J Cell Sci. 1984 Jan 1;65(1):249–63.

63. Wickham H. ggplot2: Elegant Graphics for Data Analysis. Springer-Verlag New York; 2016.

64. Bellve AR, Cavicchia JC, Millette CF, O’Brien DA, Bhatnagar YM, Dym M. Spermatogenic cells of the prepuberal mouse. Isolation and morphological characterization. Journal of Cell Biology. 1977;74(1):68–85.

65. Lai JS, Herr W. Ethidium bromide provides a simple tool for identifying genuine DNA-independent protein associations. Proc Natl Acad Sci U S A. 1992;89(15):6958–62.

66. Meers MP, Bryson TD, Henikoff JG, Henikoff S. Improved CUT&RUN chromatin profiling tools. Elife. 2019;8:e46314.

67. Corces MR, Trevino AE, Hamilton EG, Greenside PG, Sinnott-Armstrong NA, Vesuna S, et al. An improved ATAC-seq protocol reduces background and enables interrogation of frozen tissues. Nat Methods. 2017;14(10):959–62.

68. Bolger AM, Lohse M, Usadel B. Trimmomatic: A flexible trimmer for Illumina sequence data. Bioinformatics. 2014;30(15):2114–20.

69. Langmead B, Salzberg SL. Fast gapped-read alignment with Bowtie 2. Nat Methods. 2012 Mar 4;9(4):357–9.

70. Danecek P, Bonfield JK, Liddle J, Marshall J, Ohan V, Pollard MO, et al. Twelve years of SAMtools and BCFtools. Gigascience. 2021 Feb 16;10(2):1–4.

71. Ramírez F, Dündar F, Diehl S, Grüning BA, Manke T. DeepTools: A flexible platform for exploring deep-sequencing data. Nucleic Acids Res. 2014 Jul 1;42(W1):W187–W191.

72. Zhang Y, Liu T, Meyer CA, Eeckhoute J, Johnson DS, Bernstein BE, et al. Model-based analysis of ChIP-Seq (MACS). Genome Biol. 2008;9(9):r137.

73. Kim D, Pertea G, Trapnell C, Pimentel H, Kelley R, Salzberg SL. TopHat2: Accurate alignment of transcriptomes in the presence of insertions, deletions and gene fusions. Genome Biol. 2013 Apr 25;14(4):R36.

74. Anders S, Pyl PT, Huber W. HTSeq-A Python framework to work with high-throughput sequencing data. Bioinformatics. 2015 Jan 15;31(2):166–9.

75. Love MI, Huber W, Anders S. Moderated estimation of fold change and dispersion for RNA-seq data with DESeq2. Genome Biol. 2014 Dec 5;15(12):550.

76. Anand L, Rodriguez Lopez CM. ChromoMap: an R package for interactive visualization of multi-omics data and annotation of chromosomes. BMC Bioinformatics. 2022;23(1):1–9.

77. Ewels PA, Peltzer A, Fillinger S, Patel H, Alneberg J, Wilm A, et al. The nf-core framework for community-curated bioinformatics pipelines. Vol. 38, Nature Biotechnology. Nature Research; 2020. p. 276–8.

78. Lun ATL, Smyth GK. Csaw: A Bioconductor package for differential binding analysis of ChIP-seq data using sliding windows. Nucleic Acids Res. 2015;44(5):e45.

79. Heinz S, Benner C, Spann N, Bertolino E, Lin YC, Laslo P, et al. Simple Combinations of Lineage-Determining Transcription Factors Prime cis-Regulatory Elements Required for Macrophage and B Cell Identities. Mol Cell. 2010;38(4):576–89.

