## Supplemental Figures for "INO80 regulates chromatin accessibility to facilitate suppression of sex-linked gene expression during mouse spermatogenesis"

Chakraborty_Figure. S1

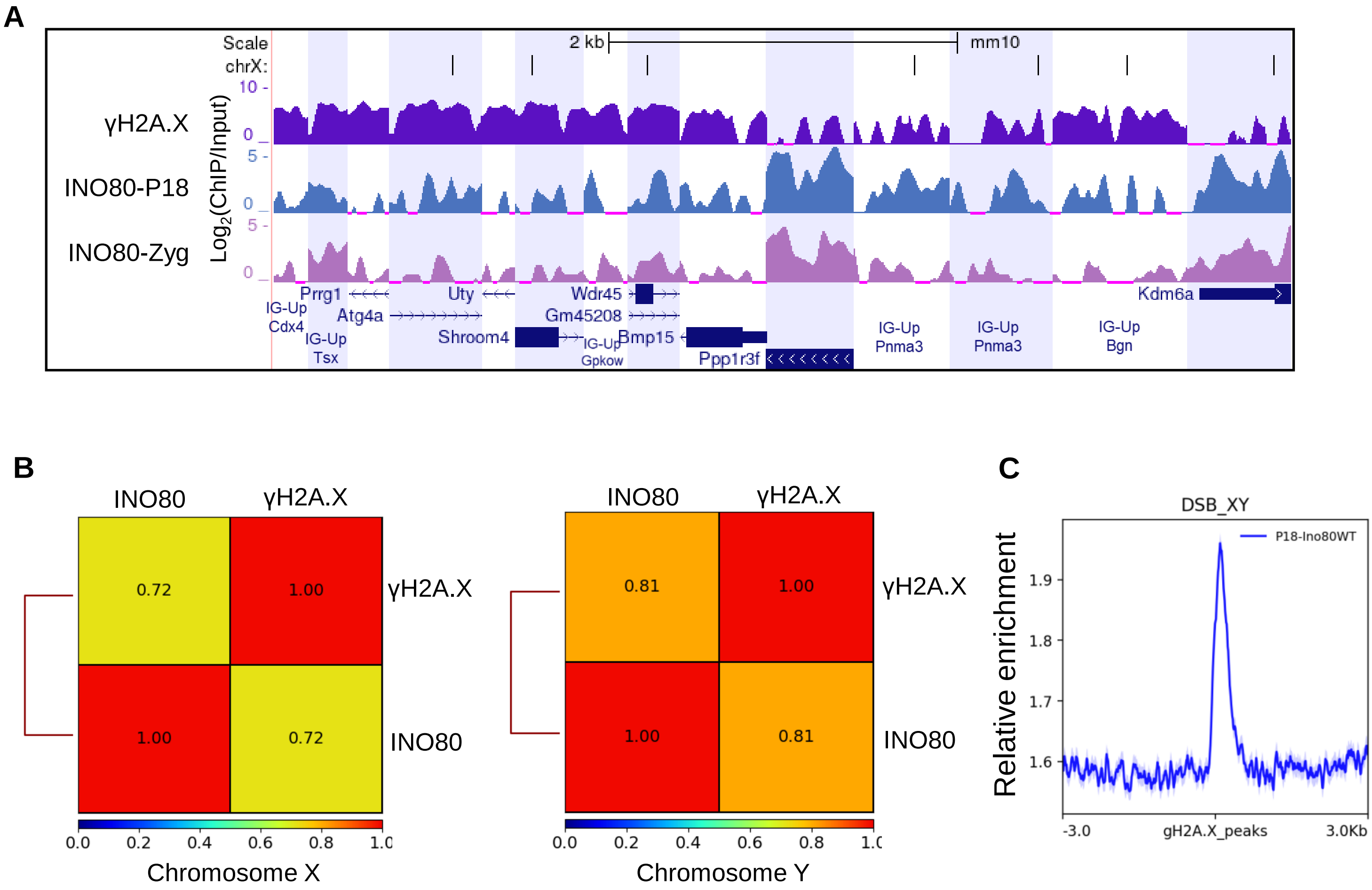

Chakraborty_Figure. S2

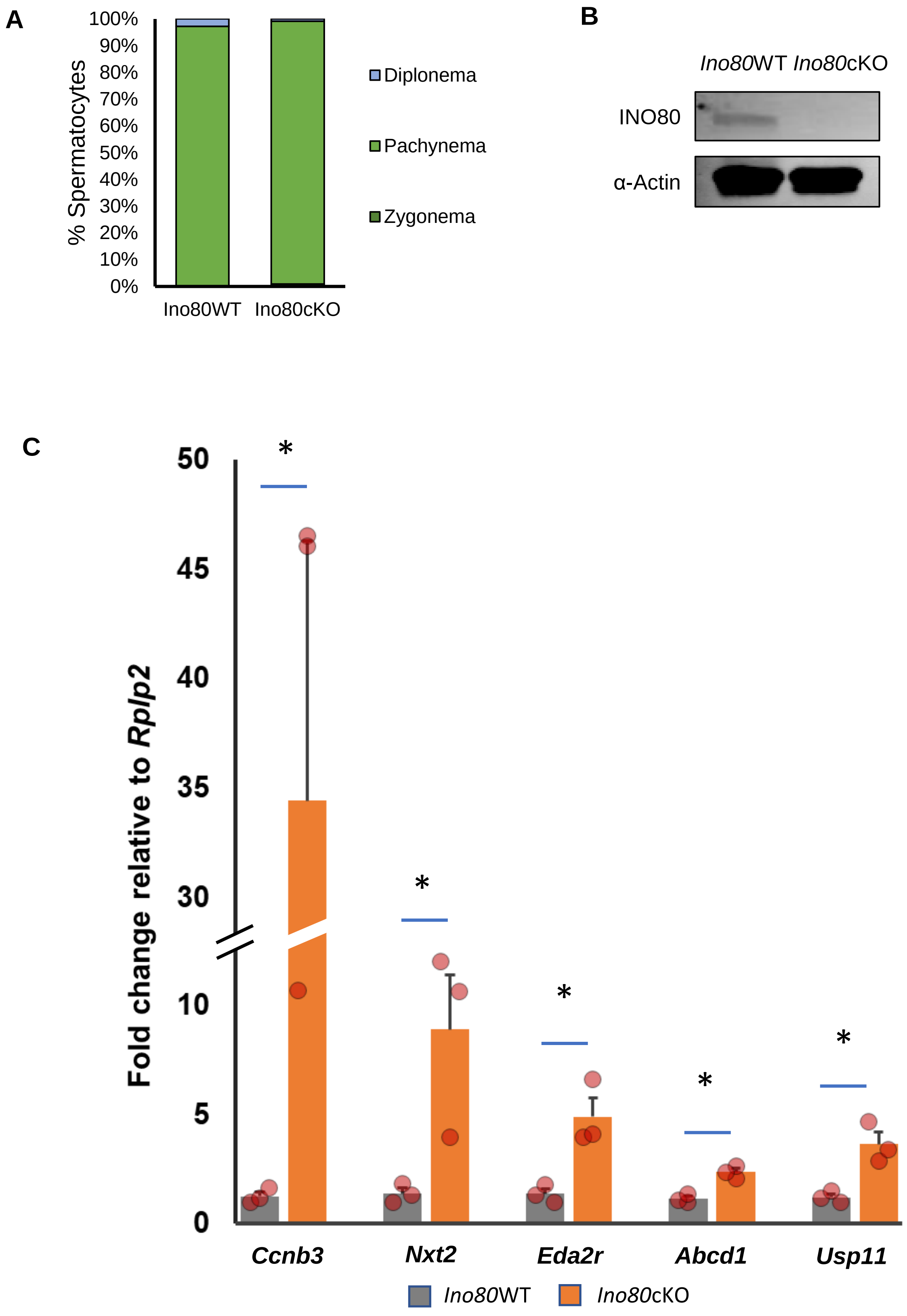

Chakraborty_Figure. S3

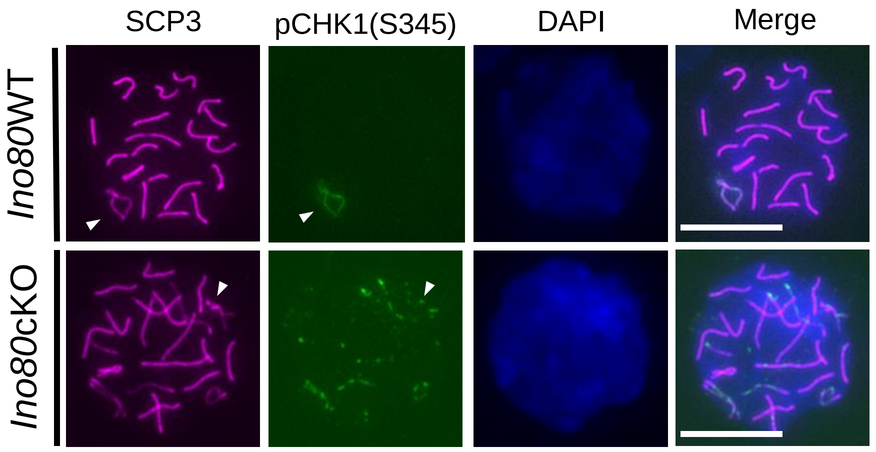

Chakraborty_Figure. S4

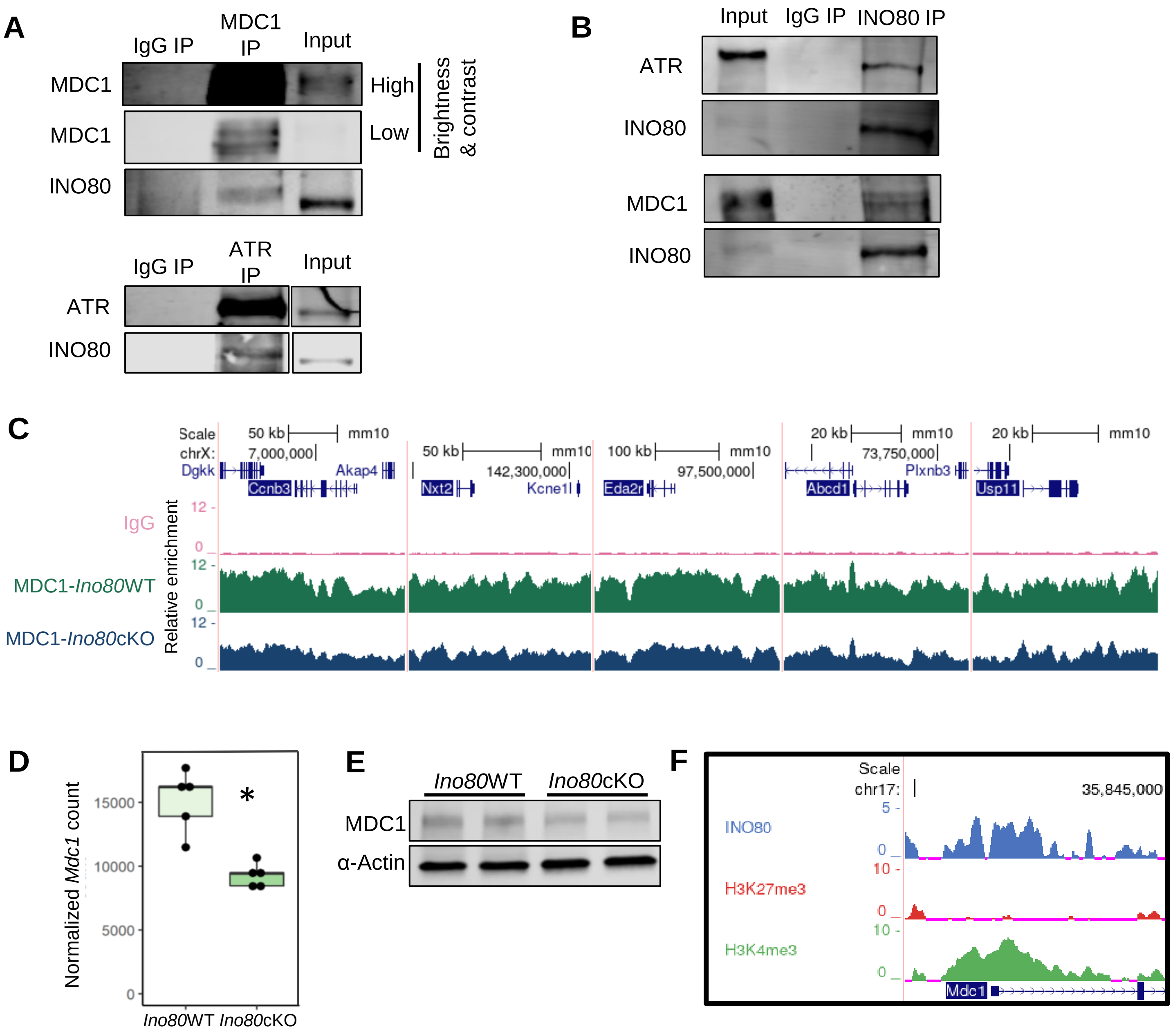

Chakraborty_Figure. S5

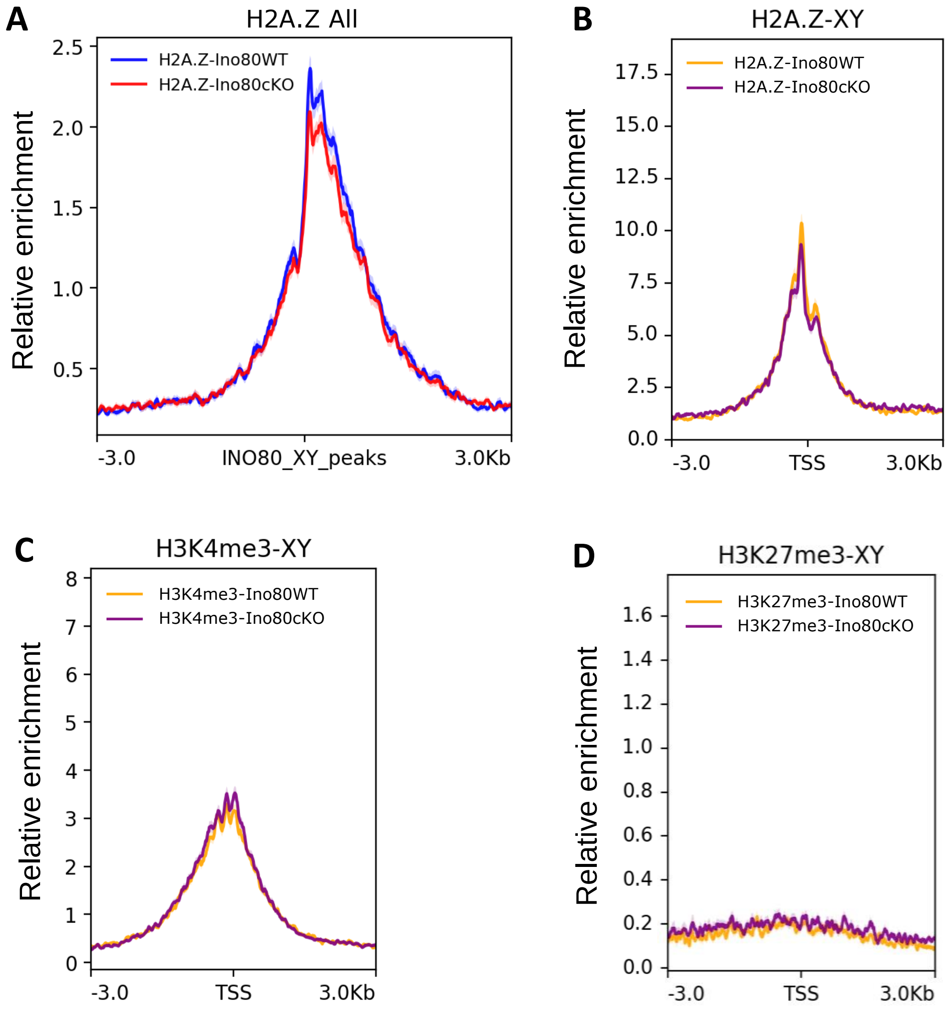

Chakraborty_Figure. S6

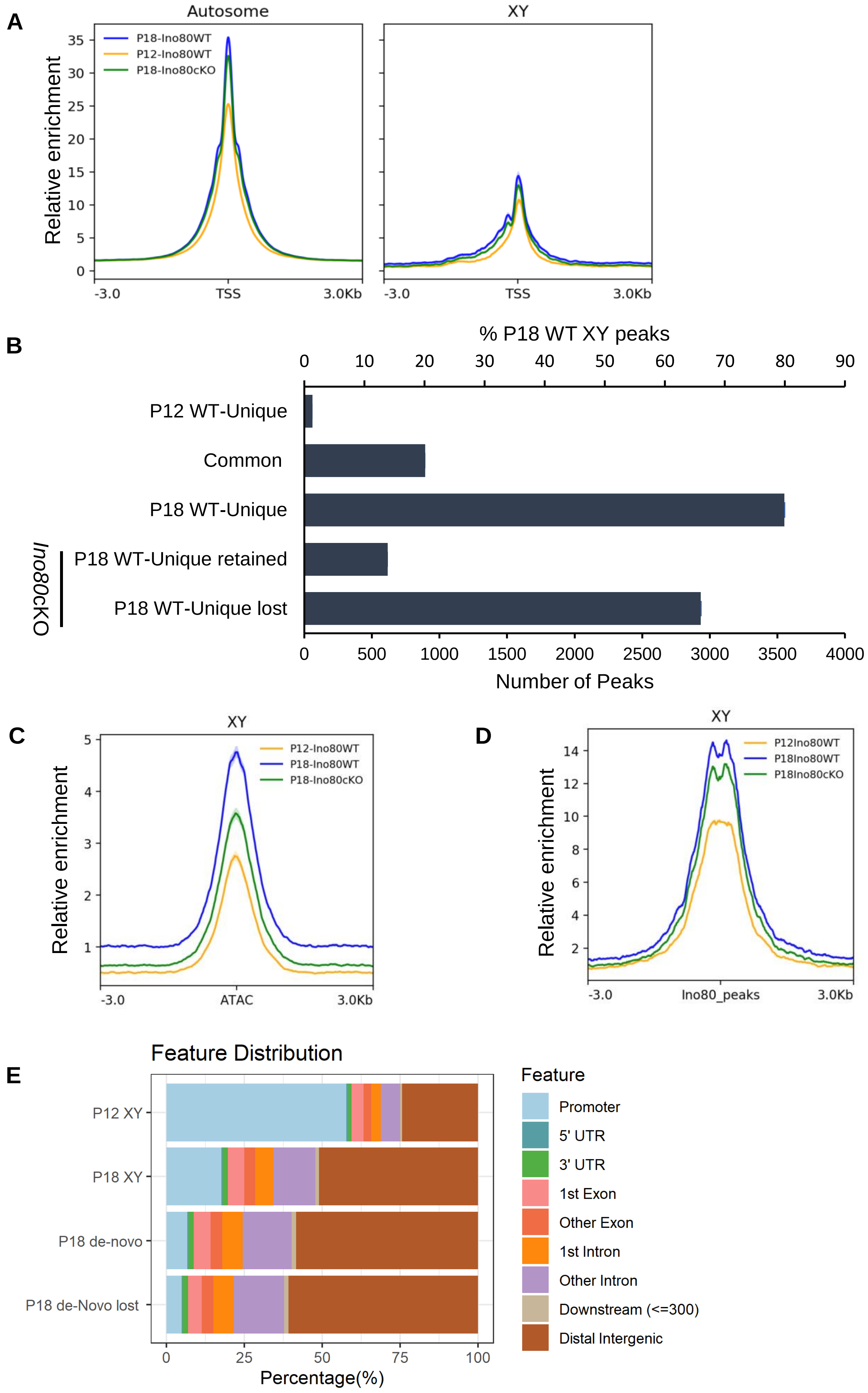
