## Supplemental Tables for "INO80 regulates chromatin accessibility to facilitate suppression of sex-linked gene expression during mouse spermatogenesis"

Table S1: Quantitative PCR primers used in this study.

| **Region/Gene** | **Primer** |
| --- | --- |
| Ccnb3 Forward | 5'-GCTCACCTCAAGCCCATTAT-3' |
| Ccnb3 Reverse | 5'-TTGACTGGTGGCTTCTCTTTAG-3' |
| Nxt2 Forward | 5'-GTAGAGCTGCCGAGGAATTT-3' |
| Nxt2 Reverse | 5'-TCCAGATTAGAGTGGCTTTGTC-3' |
| Eda2r Forward | 5'-CCAGTTGAGCTTAGTGAAGGTAG-3' |
| Eda2r Reverse | 5'-AGGAAGGCCAGAGCAAATAC -3' |
| Abcd1 Forward | 5'-TGGATGGACGACTTCGAAAC-3' |
| Abcd1 Reverse | 5'-GGCTTGGTCAGGTTGGAATA-3' |
| Usp11 Forward | 5'-GCTCGTTCAGCACAGTGATA-3' |
| Usp11 Reverse | 5'-TCCTGAGGCTCTACCAGAAA-3' |
| Rplp2 Forward | 5'-CCTAGCGCCAAAGACATCAA-3' |
| Rplp2 Reverse | 5'-GACCTTGTTGAGCCGATCAT-3' |

Table S2: Genotyping primers used in this study.

| **Allele** | **Sequence** |
| --- | --- |
| Ino80 floxed forward | 5'-GATACTTCTGCCTCCACACTTC-3' |
| Ino80 floxed reverse | 5'-CTGGCACCTTTCCAGTCTTT-3' |
| Ino80 excised forward | 5'-TGTGTAGCAACCTACAGCTA-3' |
| Ino80 excised reverse | 5'-GTTGCTGTGTCTTTGCTTTG-3' |
| Stra8Cre forward | 5'-GTGCAAGCTGAACAACAGGA-3' |
| Stra8Cre reverse | 5'-AGGGACACAGCATTGGAGTC-3' |

Table S3: Primary antibodies used in this study.

| Primary antibody | Source | Amount/Dilution |
| --- | --- | --- |
| Rabbit anti-INO80 | Abcam (ab105451) | ChIP-seq (10μg)  WB (1:2000)  IF (1:700) |
| Rabbit anti-INO80 | Novus Biologicals (NBP1-78758) | IP (10 μg)  IF (1:500) |
| Rabbit anti-ATR | Abcam (ab2905) | IP (10 μg) |
| Rabbit anti-ATR | Millipore (09-070) | WB 1:1000 |
| Goat anti-ATR | Santa Cruz (sc-1887) | IF (1:50) |
| Rabbit anti phospho-CHK1(Ser345) | Cell signaling (2348) | IF (1:200) |
| Mouse anti-MDC1 | Novus Biologicals (NBP2-12890) | WB (1:1000)  IF (1:200)  CUT&RUN (1:200) |
| Mouse anti-alpha-Actin | Santa Cruz (sc-32251) | WB 1:2000 |
| Mouse anti-SYCP3 | Abcam (ab97672) | IF 1:1000 |
| Rabbit anti-SYCP3 | Novus Biologicals (NB300-231) | IF (1:200) |
| Goat anti phosphor RNA Polymerase II (Ser2) | Active Motif (61084) | IF: (1:200) |
| Mouse anti-γH2A.X | Millipore (05-636) | IF (1:1500) |
